# Persistent co-infection mediates inter-virus and host-virus gene exchange in marine protists

**DOI:** 10.64898/2026.07.28.741200

**Authors:** Jessica Latimer, Shannon J. Sibbald, Kate Thomson, Yumi Murakoshi, Kayla Surgenor, Dudley Chung, Nikolaj Brask, Hongda Zhao, Craig McCormick, Hiroyuki Ogata, Yoshitake Takao, John M. Archibald

## Abstract

Genomics and metagenomics have transformed our understanding of the diversity of large DNA viruses infecting eukaryotic microorganisms. Using long-read sequencing, we discovered ubiquitous, co-culturing nucleocytoviruses in several thraustochytrid protists. These viruses are closely related to SmDNAV, an ‘orphan’ virus of the thraustochytrid *Sicyoidochytrium minutum*. Transcriptomics, proteomics and electron microscopy demonstrate viral gene expression and particle formation in these cultures without obvious defects in culture growth. We also found SmDNAV-type viruses associated with previously sequenced thraustochytrids. Phylogenetic analysis reveals that the SmDNAV and SmDNAV-like genomes belong to a hitherto unrecognized order of *Nucleocytoviricota*, here named “*Skiavirales*”. All fifteen identified skiavirus genomes lack numerous viral hallmark genes including DNA-dependent RNA polymerase; all but three also lack DNA polymerase family B (PolB). PolB phylogeny suggests a specific relationship between *Skiavirales* and the recently discovered mirusviruses, which belong to a different viral realm. Gene exchange between skiaviruses, co-occurring mirusviruses, and host nuclear genomes shows that persistent infection by diverse large DNA viruses provides an opportunity for virus-virus and virus-host co-evolution in thraustochytrids and, perhaps, other microbial eukaryotes.

## Introduction

Thraustochytrids are unicellular eukaryotes that are important decomposers and nutrient recyclers in the world’s oceans.^1^ Due to their high production of omega-3 polyunsaturated fatty acids (PUFAs), the metabolism of thraustochytrids has been intensively studied and they have become biotechnological workhorses in the nutritional supplement industry.^2–4^ The life cycle, ecology and evolution of these heterotrophic stramenopiles remain much less well understood. This includes their interactions with RNA and DNA viruses.

Previous studies have identified lytic viruses infecting thraustochytrids. The Aurantiochytrium RNA virus (AuRNAV; originally reported as Schizochytrium single-stranded RNA virus)^5^ possesses a genome of 9 kb^6^ and produces small (25 nm) icosahedral virus particles.^7^ The large double-stranded DNA virus of *Sicyoidochytrium minutum* (SmDNAV) was found to infect a different subset of thraustochytrids, namely two strains of the genus *Sicyoidochytrium*.^8^ During early infection, nascent viral particles accumulate in the perinuclear region, whereas nuclei are not observed in the late stages of infection.^8^ SmDNAV particles show a round morphology inside cells and a distinct “squashed ball” appearance extracellularly.^8^ The SmDNAV genome is 236 kb but lacks hallmark genes for large DNA viruses like those encoding DNA polymerase family B (PolB), DNA-directed RNA polymerase large subunit (RNAPL), DNA-directed RNA polymerase small subunit (RNAPS) and DNA topoisomerase II (TopoII).^9^ The lack of these hallmark genes in sequenced SmDNAV genomes suggests these viruses are reliant on host enzymes to support viral genome replication and gene expression. Initial phylogenetic analysis of the SmDNAV major capsid protein (MCP) suggested the virus belongs to *Nucleocytoviricota* (realm *Varidnaviria*) with similarity to *Mimiviridae* and *Phycodnaviridae*.^9^ Both AuRNAV and SmDNAV are lytic viruses with apparently very narrow, taxonomically distant host ranges and unclear ecological significance.^5^

Thraustochytrids have recently gained attention as hosts of mirusviruses (realm *Duplodnaviria*), abundant double-stranded DNA viruses whose genomes show traces of gene exchange between two distinct viral realms, *Duplodnaviria* and *Varidnaviria.*^10^ Much of the known diversity of mirusviruses comes metagenomic sequencing of marine and freshwater samples and the detection of mirusvirus genomes in a handful of eukaryotic assemblies.^10–13^ The thraustochytrid *Aurantiochytrium limacinum* has become a model organism for studying mirusviruses as cultured cells produce mature virions.^14^ *A. limacinum* has an episomal mirusvirus genome, designated AurliV-1, and an evolutionarily distinct chromosomal integrant (AurliV-2).^14,15^ Mirusvirus genes have also been detected in genomic data from other thraustochytrids such as *Hondaea fermentalgiana*^15^ and *Schizochytrium* sp. TIO01.^12^ These findings make thraustochytrids compelling hosts for the study of mirusviruses and shed new light on early descriptions of thraustochytrid viruses. In 1972, ‘herpes-type’ virus-like particles were found in a *Thraustochytrium* sp. but could not be isolated,^16^ and similar particles were seen in *Schizochytrium aggregatum* but never published.^17^ A 1979 PhD dissertation documented 150 nm virions in *Thraustochytrium aureum*.^18^

Here we used Oxford Nanopore long-read sequencing to produce high-quality genome assemblies for several well studied thraustochytrids. *Schizochytrium* sp. ATCC 20888 (also called S31), one of the best known PUFA-producing thraustochytrids and the subject of lipid yield optimization studies,^2,3,19^ was found to contain two distinct mirusvirus genomes and an SmDNAV-like viral genome, which we assign to the new order *Skiavirales*. Using transmission electron microscopy (TEM), we demonstrate the presence of both mirusvirus and skiavirus particles in *Schizochytrium* sp. 20888 cells. This is the first evidence of mirusviruses and skiavirus co-infection, both of which are characterized by long term association with cultures, constitutive viral mRNA and protein accumulation in bulk culture, and sporadic viral particle production in the absence of culture collapse. Another thraustochytrid, *Schizochytrium aggregatum* 28209, was found to be persistently infected by two skiaviruses whose genomes contain genes of mirusvirus ancestry, even though the organism lacks a mirusvirus genome. Lastly, *Thraustochytrium aureum* 34304 contains the first documented skiavirus genome with a PolB gene, phylogenetic analysis of which links *Skiavirales* to mirusviruses. Together with analysis of previously undetected viruses in publicly available thraustochytrid genomes, these findings position thraustochytrids as model systems to study skiavirus biology, their co-evolution with mirusviruses, and, more generally, the transition from lytic to persistent infection in microbial eukaryotes.

## Results

### SmDNAV-like viruses are common in, and unique to, thraustochytrid protists

Preliminary analysis of publicly available draft genomes for the thraustochytrids *Schizochytrium* sp. 20888 (www.genomes/atcc.org/products/20888), *Schizochytrium aggregatum* ATCC 28209 (Schag1, Joint Genome Institute Project ID: 402022) and *Thraustochytrium aureum* ATCC 34304 (NCBI PRJDB9722) revealed viral sequences with similarity to the genome of SmDNAV, a large DNA virus that infects *Sicyoidochytrium minutum*.^8^ Based on nuclear small subunit ribosomal RNA (SSU rRNA) phylogeny, these three thraustochytrids are not closely related to each other or to *Si. minutum*.^8,20–23^ *Schizochytrium* sp. 20888 in fact branches with *Aurantiochytrium* and *Hondaea* rather than alongside *Sc. aggregatum*.^22,23^ Given that SmDNAV completely lyses *Si. minutum* cultures within 10 days of treatment,^8,9^ the presence of SmDNAV-like sequences in *Schizochytrium* sp. 20888, *Sc. aggregatum* and *T. aureum* is intriguing. These strains have been maintained in the American Type Culture Collection (ATCC) for decades^24–26^ and studied extensively for biotechnological applications without reports of virus-induced culture collapse.^2,3,19,27,28^

To explore the possibility that SmDNAV-like viruses have gone undetected in thraustochytrids beyond *Si. minutum*, we produced chromosome-scale genomic assemblies for *Schizochytrium* sp. 20888, *Sc. aggregatum* ATCC 28209 and *T. aureum* ATCC 34304 using Oxford Nanopore’s PromethION long-read sequencing platform. A long-read genome assembly of *Schizochytrium* sp. 20888 yielded 30 nuclear contigs, one circular mitochondrial genome and three circular mapping viral contigs (Extended Data Table 1). Two viral genomes were identified as being of mirusvirus ancestry (20888-V1 and 20888-V2; Fig. 1a, b) with strong similarity to those of *A. limacinum* (Extended Data Fig. 1), while the third is a 190 kb SmDNAV-like viral genome (20888-skiaV; Fig. 1c). Our long-read re-sequencing and scaffolded assembly of *Sc. aggregatum* ATCC 28209 (Extended Data Table 1) resolved two distinct circular SmDNAV-like viral genomes (28209-skiaV1 and 28209-skiaV2; Fig. 2a, b) and the *T. aureum* assembly yielded one SmDNAV-like viral genome (34304-skiaV; Fig. 2d). As described below, comparative genomic investigation shows that SmDNAV and SmDNAV-like viruses constitute a new order of large DNA viruses, which we propose be called “*Skiavirales*”.

**Fig. 1.**
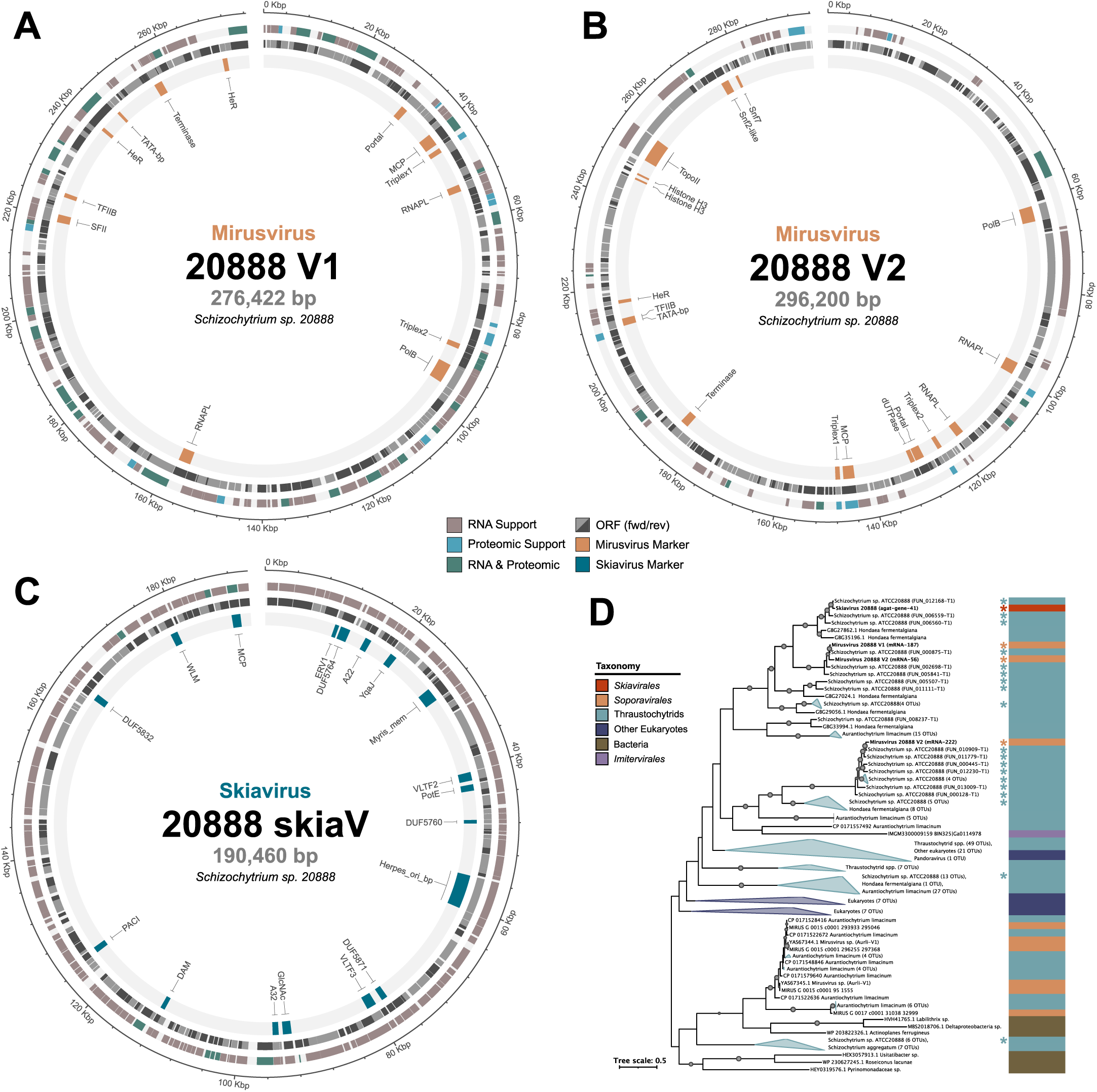
Mirusviruses and skiaviruses co-occur in the thraustochytrid *Schizochytrium* sp. 20888 and have exchanged genes with one another and their host. **a** and **b**, Physical maps of two distinct, circular mapping *Soporavirales*-type mirusvirus genomes (V1 and V2, respectively). **c**, Circular mapping skiavirus genome (20888-skiaV). **d**, Phylogenetic tree of the serine/threonine protein kinase SPS1, which suggests gene exchange between skiavirus (red), *Soporavirales* mirusviruses (orange), and thraustochytrid (host) nuclear genomes (teal). Asterisks indicate sequences from *Schizochytrium* sp. 20888.

**Fig. 2.**
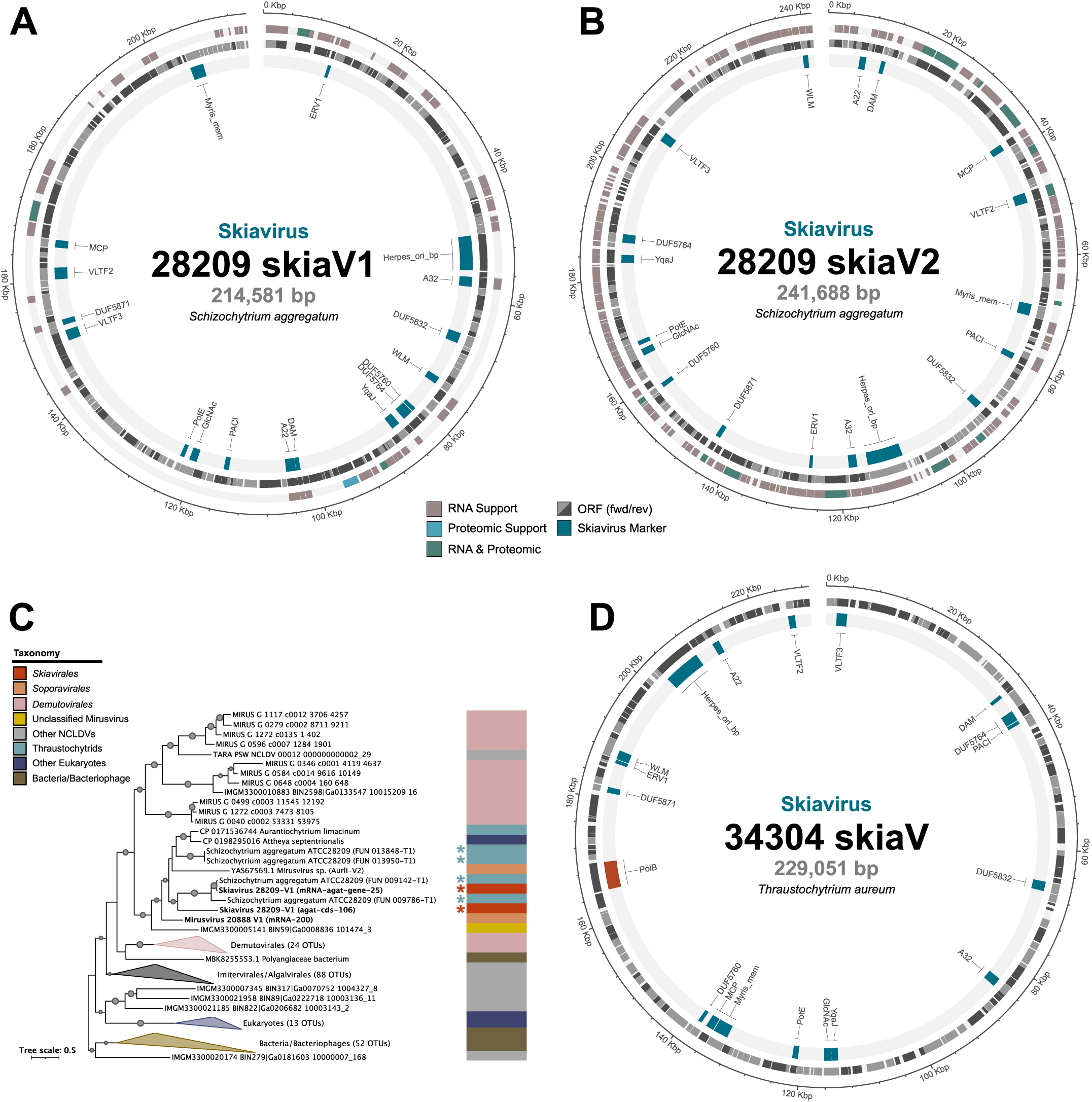
Schizochytrium aggregatum has two and Thraustochytrium aureum has one skiavirus. Physical maps of two circular mapping episomal skiavirus genomes, **a,** 28209-skiaV1 and **b,** 28209-skiaV2. **c,** Phylogenetic tree of S-adenosylmethionine-dependent methyltransferase (SAM-MTase) suggesting gene exchange between skiaviruses (red), thraustochytrid nuclear genomes (teal), as well as *Soporavirales* (orange) and *Demutovirales* (pink) mirusviruses. Asterisks indicate sequences from *Sc. aggregatum* 28209. **d,** Physical maps of circular mapping episomal skiavirus genome 34304-skiaV with DNA polymerase B (PolB) in shown in red.

Since the original SmDNAV 001 was sequenced,^9^ two other lytic strains of the virus have been isolated, sequenced and deposited in GenBank (SmDNAV Nm-Ss: LC657068 and SmDNAV Ns-Ss: LC657069). These three skiavirus genomes lack key *Nucleocytoviricota* marker genes, namely PolB, RNAPL, RNAPS and TopoII. Similarly, *Schizochytrium* sp. 20888 and *Sc. aggregatum* skiaviruses lack all these hallmark genes whereas the *T. aureum* skiavirus is one of a few rare exceptions that encodes PolB. With the uniform absence of hallmark genes in the known skiaviruses, we used alternate viral hallmark genes — MCP, A32-like packaging ATPase (A32), and virus late transcription factor 3 (VLTF3) — as bioinformatic queries to search for additional skiavirus genomes in public databases. Searches against GenBank nr, the MMETSP transcriptome database,^29^ *Tara* Oceans metagenomic data^30^ and the metavirus resource (MetaVR)^31^ identified near complete skiavirus genomes in the thraustochytrids *Parietichytrium* sp. I65-24A and *Aurantiochytrium* sp. 8W. We also Illumina sequenced and assembled the genome of *Sicyoidochytrium* sp. 09M18, and assembled raw long-read sequence data for the thraustochytrids *Aurantiochytrium* sp. TWZ-97 (NCBI SRX28266382)^32^ and *Botryochytrium* sp. S-28 (NCBI SRX19209961).^33^ These analyses yielded complete and near complete skiavirus genomes for TWZ-97 and 09M18, respectively, as well as three skiavirus genomes from *Botryochytrium* sp. S-28 (complete: S28-skiaV1 and S28-skiaV2, partial: S28-skiaV3).

Unexpectedly, we found a skiavirus genome in the dinoflagellate alga *Durusdinium trenchii* SCF082.^34^ But while the *D. trenchii* SCF082 genome assembly contains the viral sequence (contig CAXAMM010014914), the CCMP2556 reference assembly of Dougan et al.^34^ does not. The *D. trenchii* sequence data stem from a complex coral-associated microbial community, and we searched for and found thraustochytrid-type 18S rDNA sequences in the SCF082 assembly (91.54% ID to *Thraustochytriidae* sp. NK1 across 1,728 bases). Given that all other known skiavirus sequences are associated with thraustochytrids, this suggests that the *D. trenchii* sequence is in fact derived from a co-culturing thraustochytrid, not the dinoflagellate; we thus refer to it here as ‘*Durusdinium trenchii* SCF082’ skiavirus (‘SCF082’-skiaV).

We also found skiavirus gene hits in a ‘bacterial’ metagenomically assembled genome (MAG; JAAEPP010000199.1), in a contig in the genome of the ciliate *Euplotes weissei* (GCA_021440005.1), and in two environmental viral MAGs (IMGVR_UViG_3300020185_000043|Ga0206131_10000343 and IMGVR_UViG_3300045300_007864|Ga0493796_05573356). However, none of these could be resolved as *bona fide* skiavirus genomes due to low contiguity and/or sequence quality. For subsequent analyses, we focused on 15 high-quality, thraustochytrid-associated genomes: 12 identified in genomic assemblies from cultured thraustochytrids and three sequenced from SmDNAV virions (Extended Data Table 2-3, Extended Data Fig. 2-3).

### *Skiavirales*: a new order with a large pangenome

Phylogenetic analysis of three *Nucleocytoviricota* marker genes (MCP, A32, and VLTF3) reveals that all 15 complete or near complete skiavirus genomes branch robustly together as a distinct lineage of viruses sister to the medusaviruses of *Mamonoviridae*^35–38^ and the recently proposed *Manesviridae* family that includes furtivovirus,^39^ clandestinovirus,^40^ ushikuvirus,^41^ and usurpativirus^42^ (Fig. 3a, Extended Data Fig. 4). Following established protocol for *Nucleocytoviricota* taxonomy, we determined the relative evolutionary divergence (RED) for the *Skiavirales* clade to be 0.63, which falls within the range of RED scores for established *Nucleocytoviricota* orders in our trees (Fig. 3a; Order RED = 0.45-0.64). Importantly, we included the *Mriyaviricetes* in our dataset and used a different set of proteins from previous studies^43^ (MCP, A32 and VLTF3). *Skiavirales* is further subdivided in four family-level clades with RED scores of 0.77-0.88, which we refer to as SK_1-4. The sister families, *Mamonoviridae* and *Manesviridae*, have RED scores of 0.97 and 0.83, respectively.

**Fig. 3.**
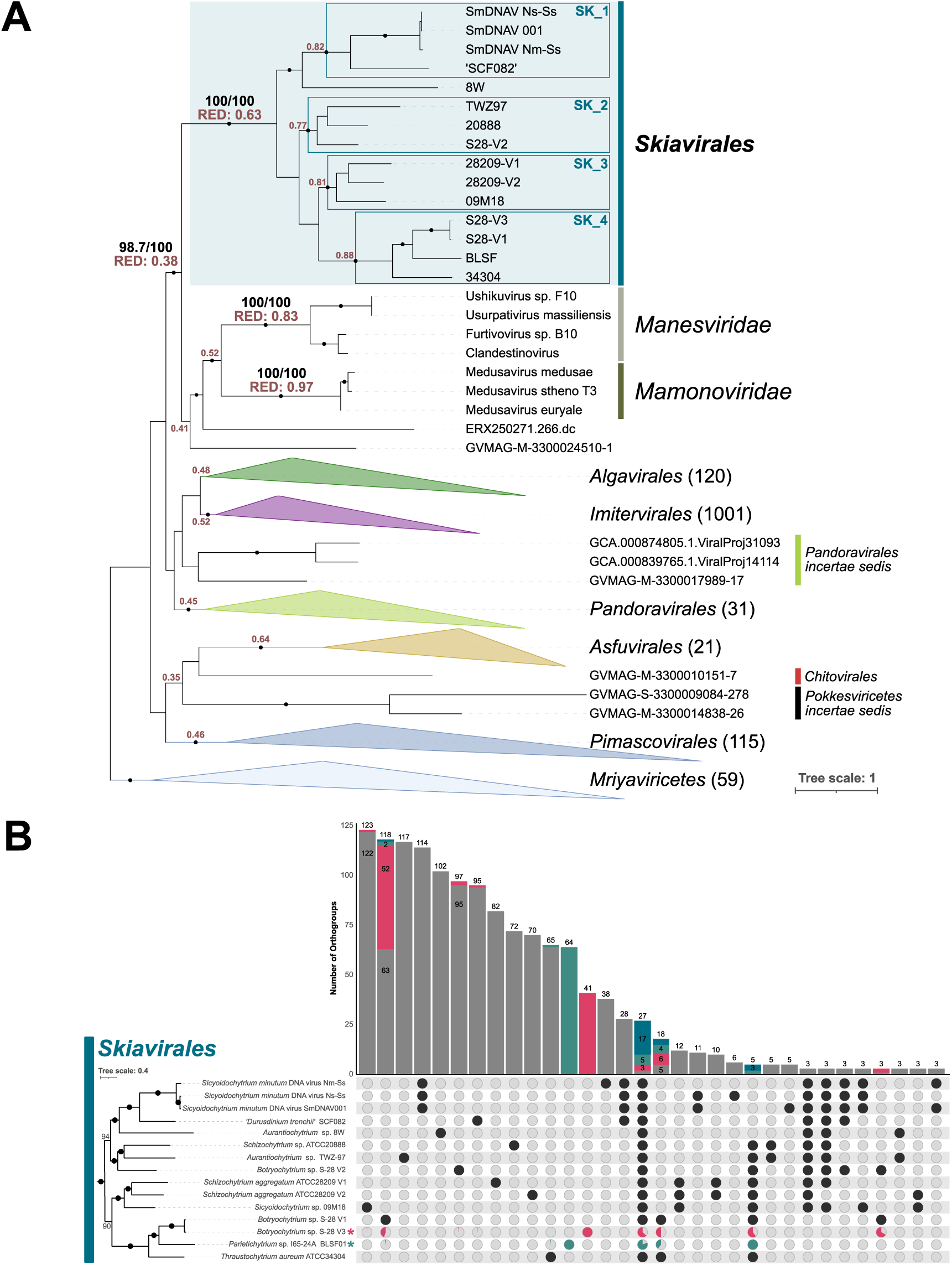
Phylogeny and pangenome of skiaviruses. **a,** Maximum likelihood phylogenetic tree of concatenated MCP-A32-VLTF3 proteins placing *Skiavirales* sister to the *Mamonoviridae* and *Manesviridae* within *Nucleocytoviricota*. The tree was inferred using IQTree under the the model selected by ModelFinder (Q.YEAST+F+I+R10, 1353 amino acids) and is rooted at *Mriyaviricetes*. Nodes with high statistical support (i.e., approximation likelihood ratios ≥ 80 and ultrafast bootstraps ≥ 95) are highlighted with dots. Relative evolutionary distance (RED) scores are shown for established *Nucleocytoviricota* orders and families, and the *Skiavirales - Manovirales* clade. **b,** Upset plot of the skiavirus pangenome showing 17 core genes for all 15 genomes. Asterisks indicate partial genomes and genes with a homolog in a partial genome are denoted within each pangenome subcategory (BLSF: teal, S28-V3: pink, or both: blue). Pangenomic data are plotted relative to the phylogenetic tree inferred from universal and near-universal single-copy orthogroups across the *Skiavirales, Mamonoviridae* and *Manesviridae*.

The node that unites *Skiavirales*, *Mamonoviridae*, *Manesviridae* and two environmental MAGs (ERX250271.266.dc and GVMAG-M-3300024510-1) branches reliably within the *Megaviricetes* (aLRT/UFboot of 98.7/100). This node has a RED of 0.38, which is lower than those of known orders and comparable to the RED score of the class *Pokke sviricetes* (RED 0.35). This suggests it is a robustly supported clade of viruses that has been greatly under-sampled in viral surveys. Using the existing gene prediction models for *Mamonoviridae* and *Manesviridae*, we identified five genes shared between all skiaviruses, mamonoviruses and manesviruses: MCP, A32, VLTF3, A22 and a YqaJ family recombinase. Except for *Medusavirus euryale*, all these viruses share a PACI endonuclease gene.

A pangenome analysis of our 15 complete or near complete skiavirus genomes identified a core genome of only 17 genes, which includes MCP, A32, VLTF3, A22, Wss1p-like metalloprotease (WLM), herpesvirus origin of replication binding protein (Herpes_ori_bp), PACI endonuclease and YqaJ-like viral recombinase domain (Fig. 3b). The skiaviruses have a large pangenome with the accessory genes dwarfing the core gene set. For example, the three skiavirus strains that lytically infect *Sicyoidochytrium* spp. cells (SmDNAV 001^8,38^, SmDNAV Nm-Ss and SmDNAV Ns-Ss; Extended Data Fig. 5) are very closely related and have a shared subset of 114 genes (Fig. 3b). Interestingly, the two persistent skiaviruses in *Sc. aggregatum* (28209-V1 and 28209-V2, both belonging to SK_3) only share 10 genes in addition to the core genes, with numerous singletons unique to each individual skiavirus; 28209-V1 has 82 and 28209-V2 has 70 singleton genes. Similarly, *Botryochytrium* sp. S28 contains three distinct skiavirus genomes, S28-skiaV1, S28-skiaV2 and a partial genome S28-skiaV3. S28-skiaV2 bears little resemblance to the other two, with 95 singleton genes and is phylogenetically assigned to a different family-level clade (SK_2). Comparatively, S28-skiaV1 and the partial skiavirus genome S28-skiaV3 both belong to SK_4 and share a subset of 55 genes that are only found to these two strains. Given this increased conservation, S28-skiaV3 likely also encodes PolB, but it was not recovered due to the partial nature of the genome. Even with this high overlap in gene content, S28-skiaV1 and S28-skiaV3 are clearly not two copies of the same skiavirus; S28-skiaV1 has 64 unique genes and S28-skiaV3 has 41.

Notably, all examined skiavirus genomes lack DNApolB, RNAPL, RNAPS and TopoII — apart from PolB in SK_4 (34304, S28-V1, S28-V3, BLSF) — which has never been documented for *Nucleocytoviricota* genomes this large. The absence of these replication- and transcription-associated genes has only recently been observed in the ‘tiny’ (35-45 kb) genomes of mriyaviruses^44^ and the mirusvirus orders *Okeanovirales* and *Styxvirales* that belong to the *Duplodnaviria*.^11^ Without these key genes, the viruses must be reliant on host machinery.

### Co-occurrence of skiavirus and mirusvirus in *Schizochytrium* sp. 20888

In *Schizochytrium* sp. 20888, two circular mirusvirus genomes were found to co-occur with skiavirus in bulk cultures. This is the second instance of persistent mirusvirus infection in a protist and the first evidence of co-infection of mirusvirus and skiavirus. The mirusvirus genomes are 276 kb (V1) and 296 kb (V2) in size and encode key informational genes (e.g., DNApolB, RNAPL) and virion morphogenesis genes (MCP, triplex 1, triplex 2, portal protein and terminase) but lack RNAPS (Fig. 1a, b; Extended Data Table 4). Note that triplex 1 was not detected with HMMs from previous studies^11^ but could be annotated by BLASTp. In a concatenated MCP-portal-terminase tree, both 20888-V1 and -V2 branch within *Soporavirales*, mirroring the phylogeny of two circular *Sopora* mirusviruses previously identified in *Schizochytrium* sp. TIO01 (Extended Data Fig. 1).^11,12^ Much like the *A. limacinum* mirusviruses^14,15^ and those of *Soporavirales*,^11,12^ the *Schizochytrium* sp. 20888 mirusvirus genomes encode their RNAPL in two distinct ORFs and the terminase is encoded by a single full-length ORF. We identified a single intron in a 20888-V1 ankyrin repeat gene with low support for a minor alternate isoform (Extended Data Fig. 6a).

RNA-seq and proteomics of bulk cell culture confirmed expression of all three *Schizochytrium* sp. 20888 viruses in nutrient rich conditions (Fig. 1a-c). The high gene density, gene expression and lack of evidence of genome erosion suggests these viruses are intact, possibly experiencing genome maintenance although the mechanism remains unclear. Additionally, TEM identified mirusvirus particles in the nucleus and cytoplasm of *Schizochytrium* sp. 20888 cells (Fig. 4b-e) though virus-containing cell sections were rare (17 of 983 observed cell sections; 1.73%). These mirusvirus particles are 176 ± 25 nm (average ± standard deviation, n = 63) in diameter and contain a punctate, electron-dense core 98 ± 11 nm (n = 111), similar to those described in *A. limacinum*.^14^ Putative skiavirus virus particles were also identified in 0.31% of observed cell sections (3 of 983 cells, two with SmDNAV-like virus particles stacked in a semi-crystalline array (Fig. 4f-i) and a cell with a single viral particle (Fig. 4j)). The putative skiavirus particles are 163 ± 14 nm (n = 117) in diameter, round, electron-dense (empty capsids can also be seen), and found in the cytoplasm (Fig. 4f-i). Notably, mirusvirus and skiavirus particles have distinct morphologies (Fig. 4e and 4j) and no cell sections obviously contained both at the same time (0 of 983 cells).

**Fig. 4.**
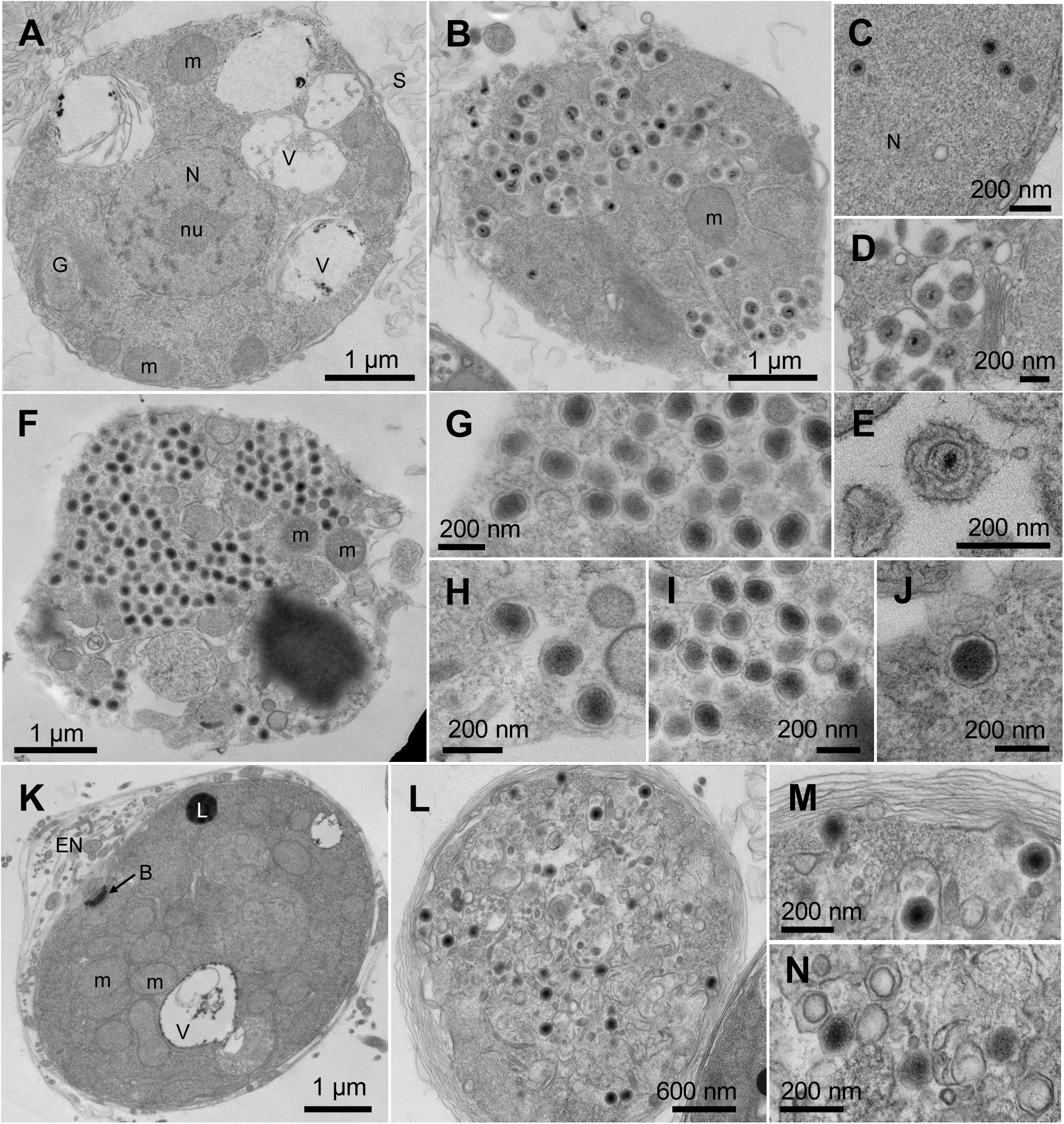
Viral particles in the thraustochytrids *Schizochytrium* sp. 20888 and *Thraustochytrium aureum*. **a,** Representative TEM image of a *Schizochytrium* sp. 20888 cell lacking obvious viral particles and **b,** a mirusvirus-producing cell. **c-e,** Close-ups showing mirusvirus particles in different sub-cellular locations. **f,** Skiavirus-producing *Schizochytrium* sp. 20888 cell with enlarged images of **g-j,** individual virus particles including empty ones. **k,** Representative TEM image of a *Thraustochytrium aureum* cell lacking obvious viral particles and **l,** a skiavirus-producing cell. Close-ups of **m,** filled and **n,** empty skiavirus particles. Abbreviations: nucleus (N), nucleolus (nu), mitochondrion (m), scales (S), Golgi apparatus (G), vacuole (V), lipid droplet (L), bothrosome (B), and ectoplasmic nets (EN).

Given the absence of key replication and transcription genes, along with a highly expressed 20888-skiaV ORFan with a strongly supported spliceosomal intron (Extended Data Fig. 6b), we infer 20888-skiaV to be reliant on nuclear host machinery for replication (note that none of the skiavirus-containing cells had a discernable cell wall or nucleus). The bulk genomic, transcriptomic and proteomic data indicate persistent association of these viruses with *Schizochytrium* sp. 20888 cultures. However, in the absence of single cell data, it is unclear if all cells vertically inherit one or both viral genomes with occasional bursts of viral production, or whether the viruses infect very few cells and is transferred horizontally between host cells.

Although mirusvirus and skiavirus particles were not observed in the same cell, their co-occurrence in long-standing bulk cultures of *Schizochytrium* sp. 20888 raises the possibility that the two viruses have exchanged genes. To investigate, we performed phylogenetic analyses of 20888-V1, -V2 and -skiaV genes. Single gene trees do not support widespread or ancient gene exchange between *Sopora* mirusviruses and the skiaviruses. We did, however, find evidence for recent gene exchange between the *Schizochytrium* sp. 20888 nuclear genome, mirusviruses, and skiavirus, including multiple copies of the serine/threonine protein kinase SPS1 (Fig. 1d) and metallophosphoesterase (MPP; Extended Data Fig. 7).

### Two skiaviruses in Schizochytrium aggregatum

As noted, the original draft short-read genome assembly of *Sc. aggregatum* ATCC 28209 (Schag1, JGI Project ID: 402022) was found to have several contigs with skiavirus marker genes. Re-sequencing with nanopore long-read technology herein resolved two distinct circular skiavirus genomes (28209-skiaV1 and -skiaV2; Fig. 2a, b; Extended Data Table 3), the first example in a single host. While neither 28209-skiaV1 or 28209-skiaV2 show obvious signs of genome erosion, only 28209-skiaV2 has notable RNA and proteomic expression, although the MCP is not detected in the cell pellet proteome (Fig. 2a, b). We did not observe skiaviruses in TEM (0 of 1,095 observed cells), consistent with previous studies of *Sc. aggregatum* that do not report on the presence of viral crystalline arrays.^27,45–47^ That said, in figures 3C, 4B, and 7A of Iwata et al.^27^ we note the presence of a few zoospores containing a ∼100 nm skiavirus-like particle. We thus hypothesize that both 28209-skiaV1 and 28209-skiaV2 persistently infect the host, but 28209-skiaV2 is more highly expressed and possibly capable of sporadic productive infections.

Given that mirusvirus-like particles were observed in *Sc. aggregatum* in 1976,^17^ we searched for, and found, mirusvirus genes in our long-read assembly of this organism, including an MCP gene. However, a complete, stand-alone mirusvirus genome was not found. Since none of the important structural mirusvirus genes (e.g., MCP) were detected by transcriptomics or proteomics, *Sc. aggregatum* may have an actively degrading mirusviral genome, one that is no longer capable of yielding productive infections.

Phylogenetic analyses revealed evidence for gene exchange between 28209-skiaV1/V2, the 28209 nuclear genome, and mirusviruses. One such gene, the S-adenosylmethionine-dependent methyltransferase (SAM-MTase) gene, is present in multiple copies in the *Sc. aggregatum* nuclear genome and in 28209-skiaV1 (Fig. 2c). Remarkably, these SAM-MTase genes are related to those found in mirusviruses: *Soporavirales* mirusviruses 20888-V1 and Aurli-V2 from *A. limacinum* branch closely to the skiaviral genes and within a larger clade of *Demutovirales* (Fig. 2c). Phylogenetic analysis also suggests that the histone H3 gene in 28209-skiaV1 and 28209-skiaV2 shares ancestry with mirusvirus homologs and host nuclear sequences, although the short length and low degree of conservation within eukaryotes make the evolutionary relationships more difficult to parse (Extended Data Fig. 8). Altogether, these data suggest frequent gene exchange between thraustochytrid hosts and their persistent viruses.

### Skiaviruses and mirusviruses share PolB ancestry

As with *Schizochytrium* sp. 20888 and *Sc. aggregatum,* our long-read genomic assembly of *T. aureum* resolved a complete circular skiavirus (34304-skiaV; Fig. 2d). However, unlike 20888 and *Sc. aggregatum*, *T. aureum* shows no trace of mirusvirus marker genes and its skiavirus genome encodes PolB. Under TEM, we observed full and empty skiavirus particles in a single cell (1 of 714 observed cell sections; 0.14%; Fig. 4l-n). The *T. aureum* 34304-skiaV particles are 130 ± 11 nm (n = 29) with a more icosahedral morphology than those in *Schizochytrium* sp. 20888 or SmDNAV 001,^8^ and may correspond to the *T. aureum* viruses described by Pollak in 1979.^18^

The presence of PolB in skiaviruses appears to be restricted to the SK_4 clade (*T. aureum* 34304-skiaV, *Botryochytrium* sp. S28-skiaV1, S28-skiaV3 and BLSF-skiaV), none of which have been previously investigated genomically. Both BLSF-skiaV (from *Parietichytrium* sp. I65-24A) and S28-skiaV3 are partial genomes; BLSF-skiaV contains a partial PolB gene and a PolB could not be found in S28-skiaV3, perhaps due to a lack of assembly contiguity. Importantly, while members of the SK_4 cluster have, or are expected to have, PolB, they appear to lack another important gene for DNA replication, the DNA sliding clamp (proliferating cell nuclear antigen, PCNA) that is found in the other 11 skiaviruses.

We took advantage of the presence of the phylogenetically informative PolB in a subset of skiavirus genomes to investigate evolutionary relationships between skiaviruses and a broader range of viruses, including mirusviruses. Intriguingly, PolB phylogenetic analysis links skiavirus clade 4 sequences to the *Soporavirales* mirusviruses with high support (Fig. 5a). The *Skiavirales* - *Soporavirales* clade is itself sister to *Demutovirales*, the other order of mirusviruses whose genomes typically contain PolB.^48^ Several skiavirus-containing thraustochytrids also have *Soporavirales* genomes with PolB (Fig. 5b, Extended Data Fig. 1), including *Parietichytrium* sp. I65-24A, whose skiaviruses and mirusviruses both have the gene (Fig. 5; BLSF and MIRUS BLSF).

**Fig. 5.**
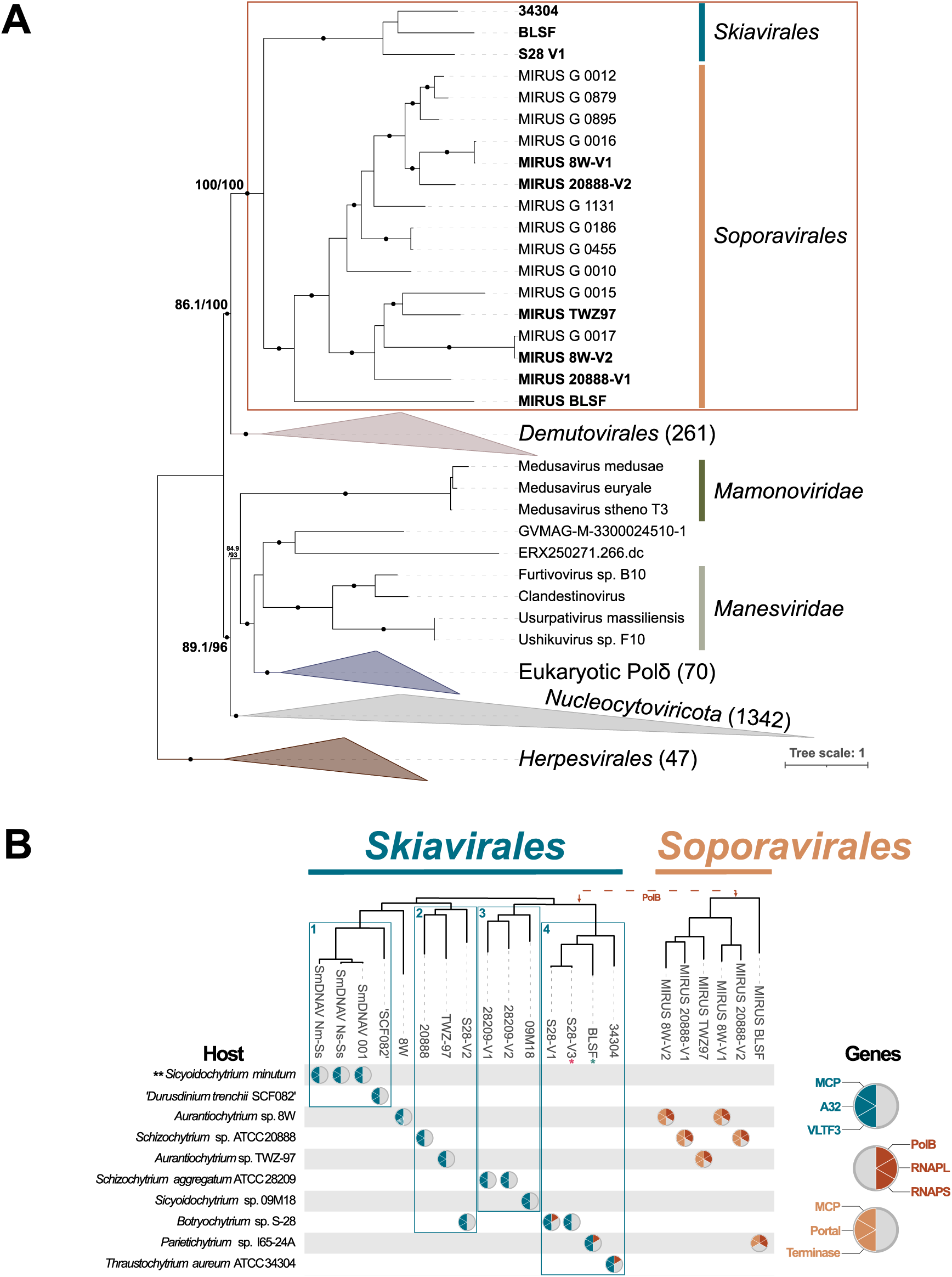
Phylogenetic tree of DNA polymerase family B showing *Skiavirales* sister to *Soporavirales* mirusviruses. **a,** IQtree maximum likelihood phylogenetic tree of PolB (1748 sequences, 2023 sites, LG+F+R10) placing the three *Skiavirales* PolB sister to *Soporavirales* PolB. The *Mamonoviridae* and *Manesviridae* branch separately from *Skiavirales* and the rest of *Nucleocytoviricota*. The tree is rooted at *Herpesvirales*. Nodes with high statistical support (i.e., approximation likelihood ratios ≥ 80 and ultrafast bootstraps ≥ 95) are highlighted with dots. **b,** Summary of skia and mirusviruses detected across hosts with skiavirus marker genes (blue: MCP, A32 and VLTF3), mirusvirus marker genes (orange: MCP, Portal and Terminase), and replication genes (red: PolB, RNAPL and RNAPS) indicated. Asterisks indicate partial genomes (BLSF and S28-V3). Double asterisks denote *Sicyoidochytrium minutum* which is not a persistently infected host as it is lysed by all three strains of *Sicyoidochytrium minutum* DNA virus (SmDNAV).

Notably, *Mamonoviridae* and *Manesviridae* branch separately from *Skiavirales* and basal to eukaryotic Polδ rather than with the rest of *Nucleocytoviricota* (Fig. 5a), which is where medusaviruses branch in previous PolB phylogenies.^35,49^ This suggests that the specific relationship between skiaviruses and mirusviruses observed here does not extend to the *Mamonoviridae* and *Manesviridae,* and is more likely the result of recent virus-virus gene sharing within a thraustochytrid host. This hypothesis is supported by phylogenetic analysis of DNA adenine methyltransferase (DAM), a core skiavirus gene that is specifically related to DAM homologs found in members of the *Demutovirales* (Extended Data Fig. 9). These data suggest that gene exchange between different realms of DNA viruses has occurred within thraustochytrid hosts.

## Discussion

Our results support the existence of skiaviruses, a novel group of *Nucleocytoviricota* with a host range apparently restricted to thraustochytrids and a striking lack of informational genes.^44,50^ We propose the order level name *Skiavirales* (*skia*: Greek for ‘shadow’ or ‘shade’ and used to describe ethereal spirits of the underworld) inspired by the electron dense viral particles observed by TEM and the fact that the viruses ‘haunt’ their hosts over long timescales, in the lab and presumably in nature. This naming scheme conforms to the etymology of related viral families *Manesviridae* (*manes*: ancient Roman chthonic deities or spirits of the dead) and *Mamonoviridae* (*mamono* (魔物): Japanese for ‘monster’ and *medusa* genus nomenclature inspired by the Gorgons of Greek mythology).

Phylogenetic analysis suggests that *Skiavirales* are sister to *Mamonoviridae* and *Manesviridae* (Fig. 3a) (two MAGs (ERX250271.266.dc and GVMAG-M-3300024510-1) also branch in this region of the tree, one from fresh water and the other from palm oil mill effluent^43^). A recent study proposed that these two families constitute a novel order within the phylum *Nucleocytoviricota*^39^ which has been tentatively assigned the name *Manovirales.*^51^ The robust support for the *Skiavirales - Manovirales* clade is particularly notable, with a RED score comparable to existing classes (e.g., *Pokkesviricetes* RED*=*0.35). This clade falls reliably within the *Megaviricetes*, and given the absence of key marker genes in skiavirus genomes, we are only able to construct a concatenated tree with some of the shorter and less informative *Nucleocytoviricota* markers (MCP, A32 and VLTF3); caution is thus warranted when interpreting RED scores.

Within the *Skiavirales - Manovirales* clade, there is substantial variation in genome size and coding capacity. *Manesviridae* genomes are 560-670 kb in size,^39–42^ *Mamonoviridae* genomes are 369-381 kb,^35–38^ and the genomes of *Skiavirales* described here are 190-252 kb. Interestingly, the three strains of skiaviruses (SmDNAVs) that lytically infect *Sicyoidochytrium* spp. cells (SmDNAV 001, SmDNAV Nm-Ss and SmDNAV Ns-Ss) are predicted to have linear genomes like the *Mamonoviridae*^35–38^ and *Manesviridae*,^39–42^ whereas the long-read assembled skiaviral genomes presented in this work are all circular mapping (i.e., 20888-skiaV, 28209-skiaV1, 28209-skiaV2, 34304-skiaV, S28-skiaV1, S28-skiaV2, TWZ97-skiaV and 8W-skiaV). The original SmDNAV genomes were all sequenced from concentrated viral fractions, which may reflect differences in genome architecture when the viral genome is replicating in the host compared to when it is packaged into virions.

A mere five genes were found to be shared between *Skiavirales* and *Manovirales* (MCP, A32, VLTF3, A22 and YqaJ), a viral lineage that appears to have undergone substantial genome reduction. TopoII is absent in both orders, RNAPL and RNAPS were lost in both *Skiavirales* and *Mamonoviridae*, and both orders appear to either lack PolB or encode a divergent copy (Fig. 5a). The medusaviruses of *Mamonoviridae* are known to encode a highly divergent copy of PolB that branches basally to *Nucleocytoviricota* PolB and the eukaryotic DNA polymerase delta,^35,43,49^ and our study strongly suggests that the other *Manovirales* also possess this divergent homolog (Fig. 5a). The nested placement of *Mamonoviridae*, *Manesviridae* and eukaryotic DNA polymerase delta (EukPolδ) within a broader viral clade of *Nucleocytoviricota* and *Mirusviricota* could be explained by (i) acquisition of PolB from an early eukaryotic lineage or (ii) a virus-to-eukaryote transfer before the diversification of extant eukaryotes. Our data suggest that the common ancestor of the *Skiavirales - Manovirales* clade could have been the donor or recipient — namely the lack of PolB in most skiaviral genomes analyzed thus far and the distinct branching position of the few PolB-containing skiaviruses separately from *Manovirales*, which may correspond to a distinct acquisition of a mirusvirus-like PolB (Fig. 5a). The unusual PolB ancestry of the *Skiavirales - Manovirales* clade makes them a potentially useful clade of study to elucidate the origins of giant viruses relative to their eukaryotic hosts.

Members of the *Skiavirales - Manovirales* clade infect diverse protists; the *Mamonoviridae* infect *Acanthamoeba castellanii*,^35–37^ the *Manesviridae* infect *Vermamoeba vermiformis*,^39–42^ and the *Skiavirales* infect various thraustochytrid species. It is unclear if the diverse hosts and genome sizes stem from an ancient diversification of the viruses alongside their eukaryotic hosts (Bigyra and Amoebozoa), or if there was a more recent host switching accompanied by substantial genome expansion or reduction. More extensive sampling will be needed to resolve this issue. In any case, the infection biology of skiaviruses is remarkable. *Schizochytrium* sp. 20888, *Sc. aggregatum* and *T. aureum* have been cultured for more than 30 years^24–26^; we have not observed virus-related culture crashes and are not aware of reports to that effect. At the same time, our TEM data show that *Schizochytrium* sp. 20888 and *T. aureum* can produce skiavirus particles under nutrient rich growth conditions. While we did not observe skiavirus particles in *Sc. aggregatum*, a previous study by Iwata et al. revealed some candidate skiavirus-like particles under TEM, albeit smaller (∼100 nm) than those observed in *Schizochytrium* sp. 20888.^27^ Thus while skiaviruses do not obviously cause an acute infection of entire cultures (outside of *Sicyoidochytrium* spp.), our data cannot speak to whether the mature viral particles cause lysis of individual cells when produced *en masse*. Furthermore, our results are based on bulk culture biomass and thus cannot be used to assess the prevalence of skiavirus genome copy number in each cell; we do not yet know if all *Schizochytrium* sp. 20888 and *T. aureum* cells have the genome and capacity to generate viral particles, or if the genome is absent from most cells and chronic, low-level productive viral replication serves to ‘re-seed’ the culture. Single cell genomics and transcriptomics will be required to distinguish between these possibilities.

Evidence for low-level persistence of large DNA viruses in microbial eukaryotes is growing. Recent studies of giant endogenous viral elements in the green alga *Chlamydomonas reinhardtii*^52^ and multiple species of the brown alga *Ectocarpus*^53^ show how giant viruses use latency as a viral infection strategy. While the *Manesviridae* have slower infection cycles with less obviously acute cytopathic effects than faustovirus, their genomes do not appear to be stably transmitted between hosts during host cultures in the laboratory.^40–42^ However, the skiaviruses described herein appear to be intrinsic to their host cultures. Indeed, the persistent *Schizochytrium* sp. 20888, *Sc. aggregatum* and *T. aureum* skiaviruses were only detected by bulk genome sequencing and confirmed with extensive TEM in *Schizochytrium* sp. 20888 and *T. aureum*. More research is needed to investigate the prevalence and biological significance of persistently associated viruses with protist cultures.

Mirusviruses were initially proposed to be chimeric viruses with virion morphogenesis module genes like those of *Duplodnaviria* (including herpesviruses) and an informational module of *Varidnaviria* ancestry.^10^ But while the first mirusviruses to be analyzed belong to the order *Demutovirales*, other orders, such as *Okeanovirales* and *Styxvirales*, lack some or all of the informational module genes.^11^ How might this be explained? We propose that skiaviruses are a missing link between the *Nucleocytoviricota* and mirusviruses, and that the thraustochytrids have served as the host facilitating this exchange.

We have shown that *Schizochytrium* sp. 20888 contains multiple persistent infections — two mirusvirus and one skiavirus — and the presence of multiple copies of SPS1 and MPP in nuclear and both viral genomes (Fig. 1d, Extended Data Fig. 7) broadly supports the hypothesis that co-infection serves as an opportunity for virus-virus and virus-host gene exchange. Similarly, while *Sc. aggregatum* has two skiaviruses and no obvious mirusvirus genomes, its skiaviral genomes have numerous genes of apparent mirusvirus origin (more specifically, *Soporavirales)* (Fig. 2c and Extended Data Fig. 8). This suggests the past presence of a mirusvirus genome that left a ‘footprint’ of *Soporavirales* genes in the host nuclear genome and that of co-infecting skiaviruses. The existence of a once active mirusvirus in *Sc. aggregatum* is corroborated by a single TEM image of a cell with a mirusvirus-like particle from a book published in 1976,^17^ a virus that has since been lost.

Interestingly, while the *T. aureum* genome presented herein does not have an obvious mirusvirus genetic footprint, it provides the most direct connection to the *Soporavirales*. In PolB phylogenies, the *T. aureum* 34304-skiaV branches with the other SK_4 sequences, sister to the *Soporavirales* within a larger clade of mirusvirus including *Demutovirales* (Fig. 5a). This suggests that the few skiaviruses with PolB have a mirusvirus-like PolB rather than one specifically related to the *Manovirales* or the rest of *Nucleocytoviricota.* Curiously, only one of the several thraustochytrids hosts that contain both skiaviruses and mirusviruses have PolB in each viral genome (Fig. 5b). It thus remains unclear if this PolB acquisition from mirusviruses occurred in the ancestor of SK_4 or in the ancestor of all skiaviruses and was subsequently lost in other lineages. A single acquisition in an ancestral SK_4 is most parsimonious, but the latter scenario of reductive evolution is an interesting parallel to the current models for mirusvirus evolution.

The four currently recognized mirusvirus orders have a patchy distribution of PolB, RNAPL and RNAPS, a pattern that is hypothesized to be the result of either a cytoplasm-to-nucleus viral transition with substantial gene loss and intron gain, or a nucleus-to-cytoplasm transition coupled with gene gain and intron loss.^48^ The PolB-containing *Demutovirales* are predicted to be produced in the nucleus; they have highly complete DNA replication and transcription modules, but are related to Pol-B lacking families that are not obviously nuclear or cytoplasmic. The *Demutovirales* are thus potentially an evolutionary intermediate.^48^ In *Soporavirales* of thraustochytrids, RNAPS is consistently absent (Figs. 1a,1b and 5b) and their genes harbour few introns (Extended Data Fig. 6a), yet the *Skiavirales* lack many replication and transcription genes with notable introns (Extended Data Fig. 6b). This pattern is consistent with the ’evolutionary trap’ model^48^ of gene loss and spread of introns in *Soporavirales* and *Skiavirales*. However, this model of gene loss does not consider multiple viruses in a single host system.

We propose that persistent co-infection of thraustochytrids by mirusviruses and skiaviruses has facilitated frequent virus-virus gene exchange and genome reduction. We have shown proteomic expression of a PolB-lacking skiavirus in *Sc. aggregatum* (Fig. 2a and b), the production of PolB-lacking skiavirus particles in *Schizochytrium* sp. 20888 (which has two soporaviruses; Fig. 1a-c, Fig. 4), and PolB-containing skiavirus particles in *T. aureum* (Fig. 4). Together these data suggest that co-infection is not necessary for replication of both viruses; nor does it preclude it. The PolB-lacking skiaviruses encode the replication DNA sliding clamp PCNA, which both PolB-containing skiaviruses and soporaviruses lack. Such complementary gene sets could conceivably offset the fitness cost of gene loss, leading to dependency on the host and co-infecting viruses. The evolutionary dynamics of skiaviruses and mirusviruses shown here make a compelling case for virus-to-virus gene exchange in addition to virus-to-host and/or host-to-virus gene transfer at the origin of giant viruses. Such transfers represent an unexplored aspect of virus research, one with important implications for host-virus co-evolution.

## METHODS

### Transmission Electron Microscopy

Cells were pelleted (3220 x g for 10 min) and fixed in 2.5% glutaraldehyde prepared in 1.5X PHEM buffer^54^ and 9% sucrose (*Sc. aggregatum* and *T. aureum*) or 3X PHEM (*Schizochytrium* sp. 20888) at 4°C for 90 hrs. Samples were rinsed three times with 0.1 M sodium cacodylate buffer, fixed for two hrs with 1% osmium tetroxide, then rinsed with distilled water. Samples were stained overnight at 4°C with 0.25% uranyl acetate, dehydrated using a graduated series of acetone (50%, 70% twice, 95% twice, 100% twice), embedded with Epon Araldite resin (3:1 dried 100% acetone to resin for 3 hours, 1:3 dried 100% acetone to resin overnight, 2x 3 hrs 100% resin), and cured at 60°C for 48 hrs. Thin (80-100 nm) sections, obtained using a Reichert-Jung Ultracut E ultramicrotome, were placed on 300 mesh copper grids and further stained with 2% aqueous uranyl acetate and lead citrate. Samples were imaged with an AMT Nanosprint 15L-Mk2 camera attached to a FEI Tecnai-12 microscope at 80kV.

### DNA Extraction, Genome Sequencing and Assembly

For long read Oxford Nanopore Technologies (ONT) sequencing, *Sc. aggregatum, Schizochytrium* sp. 20888 and *T. aureum* were cultured for three days in 790 By+ (1 mg/mL yeast extract, 1 mg/mL peptone, 5 mg/mL D+ glucose) in artificial seawater (ASW; 24.72 mg/mL NaCl, 0.67 mg/mL KCl, 1.364 mg/mL CaCl_2_ • 2H_2_O, 4.66 mg/mL MgCl_2_ • 6H_2_O, 6.29 mg/mL MgSO_2_ • 7H_2_O, 0.18 mg/mL NaHCO_3_). Genomic DNA (gDNA) was extracted using QIAGEN’s Deasy® Blood & Tissue Kit Quick-Start Protocol with an overnight Proteinase K digestion. Agarose gel electrophoresis (1%) was used to visually assess and confirm the integrity of high molecular weight (20+ kb) DNA. The DNA quantity was assessed using a Qubit 2.0 Fluorometer (ThermoFisher Scientific) with the dsDNA broad range assay kit. Libraries were prepared using the ONT ligation sequencing kit (SQK-LSK114), and *Sc. aggregatum* and *T. aureum* were run on a PromethION flow cell (FLO-PRO114M) while *Schizochytrium* sp. 20888 was sequenced using a minION flow cell (FLO-MIN114). Base calling was performed with Dorado (v1.0.2+c758d2f6) using super accurate model v5.0.0, 400 bps and trimming adaptors. Middle adaptors were removed by Porechop *ab initio* (v0.5.0) and data filtered using Filtlong (v0.2.1) for genome assembly with Flye (v2.9.2-b1786) in isolate mode (*Schizochytrium* sp. 20888 and *T. aureum*) and metagenome mode (*Sc. aggregatum*). The *Sc. aggregatum* genome was then scaffolded using longstitch (ntLink-arks) (v1.0.5)^55^ and gaps were closed using TGS-GapCloser using default parameters (v1.2.1).^56^

The fastq long read ONT data from previous studies on *Aurantiochytrium* sp. TWZ-97 ^32^ and *Botryochytrium* sp. S-28 ^33^ was downloaded from NCBI (SRR33000245 and SRR23265714) filtered using Filtlong (v0.2.1) and assembled with Flye (v2.9.2-b1786) in isolate mode.

*Sicyoidochytrium* sp. 09M18 was cultured in dGPY for 3 days; DNA was extracted using DNA Suisui-F (Rizo, Inc., Tsukuba, Japan) according to the manufacturer’s instructions. Illumina DNBSEQ-G400 (150bp pair-end) sequencing followed by adaptor trimming and read filtering (minlength=40, minavgquality=15) with BBduk (v39.01) was performed by bitBiome, Inc. (Tokyo, Japan). Short read assembly was done using Spades (v3.15.5) in isolate mode.

### Gene Prediction and Annotation

Viral gene prediction was performed using GeneMarkS in Virus mode (v4.28)^57^ and Prodigal (v2.6.3)^58^ with translation table 11. The initial gene predictions were compared and merged using AGAT (v1.4.0), followed by manual curation in Geneious Prime (https://www.geneious.com) with RNA sequencing support mapped with HISAT2 (v2.2.1)^59^ to identify well-supported introns, defined as those with a minimum 10 reads spanning the intron and reduced RNA-seq coverage across the intron. RNA sequencing data was obtained from the Marine Microbial Eukaryote Transcriptome Sequencing Project^29^ for *Sc. aggregatum* ATCC 28209 and from a prior study of *Schizochytrium* sp. 20888.^3^ Nuclear gene prediction was done with the RNA-seq enabled funannotate pipeline.^60^ Functional annotation of viral genes were assigned using interproscan (v5.59-91.0), BLASTp to the nr database (e-value ≤ 1e-3), CDD database, and pHMMer (version hmmer3.1b2) using the REFSEQ viral database (release version 2025/05/05). RNA expression was computed for each gene in transcripts per million (TPM). tRNA genes were predicted with both ARAGORN (v1.2.41)^61^ and tRNAscan-SE (v2.0.12) using the maximum sensitivity mode.^62^

### Orthogroup Inference

*Skiavirales* orthogroups were inferred with OrthoFinder (v 2.5.5)^63^ using blast for sequence search, a relaxed MCL inflation parameter (I=1.3) and multiple sequence alignment using MAFFT. Predicted protein-coding gene sets for *Manesviridae* (4 species) and *Mamonoviridae* (3 species) were used as an outgroup for orthogroup inference. Initially inferred orthogroups were manually curated using blast-based approaches. For species tree inference, the MAFFT-based alignment of universal/near-universal single-copy orthogroups found within *Skiavirales + Manovirales* by OrthoFinder was trimmed using TrimAl (v1.4.rev15)^64^ with a 20% gap threshold. The resulting species tree was inferred using 15,317 parsimony-informative sites in IQ-TREE2 (v 2.3.6)^65^ with the model LG+C10+F+R10 with 1000 UFboot2.^66^

### Phylogenetic Trees

All predicted protein-coding genes in *Schizochytrium* sp. 20888 and *Schizochytrium aggregatum* V1/V2 skiaviruses were queried against the NCBI nr, MMETSP transcriptome and VFAM (release233) databases as well as against a custom database consisting of Aylward et al. (2021)^43^ and Vasquez et al. (2026)^67^ viral datasets, Mirusviruses annotated in *Schizochytrium* sp. 20888 and host predicted proteins (see above) using blastP (v 2.15.0) with an e-value cut-off of 1e-5. Sequences were aligned using MAFFT-linsi^68^ and trimmed using TrimAl (v1.4.rev15)^64^ with a 20% gap threshold for alignment sites and a custom python script to remove sequences consisting of >70% gaps. Single gene trees were inferred using IQ-TREE2 (v 2.3.6)^65^ using the model selected by ModelFinder using BIC criteria^69^ and with 1000 UFboot2.^66^ All other predicted protein-coding genes in *Skiavirales* strains were queried similarly using blastP to determine the proportion of ORFans in each genome.

For multi-gene trees, the MCP, A32 and VLTF3 protein datasets from the published giant virus framework^43^ were supplemented with *Mriyaviricetes*,^44^ *Mamonoviridae*,^37,38^ *Manesviridae*,^39–42^ and the *Skiavirales* described herein. For mirusviruses, both *Schizochytrium* sp. 20888 mirusviruses were added to the Medvedeva et al. (2025)^11^ dataset. Individual gene alignments were generated using MAFFT-linsi^68^ and trimmed using TrimAl (v1.4.rev15)^64^ with a 10% gap threshold for alignment sites. Single gene trees were inferred using IQ-TREE3 (v3.0.1)^70^ using the model selected by ModelFinder using BIC criteria^69^ and with 1000 SH-aLRT^71^ and 1000 UFboot2.^66^ Multi-gene trees were inferred with the same methods using a concatenation of trimmed single gene alignments. Relative evolutionary distance (RED) scores were computed for the *Nucleocytoviricota* multi-gene tree by applying the ‘get_reds’ function of the castor R package.

We collected herpesvirus PolB and eukaryotic polymerase B delta sequences,^49^ *Nucleocytoviricota* PolB sequences,^43^ and *Mirusvirocota* PolB sequences^48^ for our PolB phylogeny. These sequences were also supplemented with *Mamonoviridae*,^37,38^ *Manesviridae*,^39–42^ and the *Skiavirales* and *Soporavirales* PolB sequences described herein. Additionally, we identified Polδ sequences from the hosts of *Skiavirales* + *Manovirales*, i.e., our thraustochytrids, *Acanthamoeba castellanii*, and *Vermamoeba vermiformis*. Sequences longer than 650 amino acids were aligned with MAFFT-fftnsi,^68^ trimmed using TrimAl (v1.4.rev15)^64^ with a 10% gap threshold for alignment sites and a gene tree was inferred using IQ-TREE3 (v3.0.1)^70^ with the model LG+F+R10 and 1000 SH-aLRT^71^ and 1000 UFboot2.

### Proteomics

One-day-old pre-seed cultures of *Sc. aggregatum* and *Schizochytrium* sp. 20888 (grown in 790 By+) were used to start cultures grown for three days at 27°C and spun down at 3220 g for 10 minutes to collect cell pellets. A whole cell data-independent acquisition (DIA) proteomics experiment was performed for *Sc. aggregatum,* and a data-dependent acquisition (DDA) proteomics experiment was conducted for *Schizochytrium* sp. 20888.

Our protocol was adapted from Hughes et al.^72^, as implemented in Chung et al.^14^ Briefly, samples were purified using SP3 beads and digested on the beads using trypsin according to Hughes et al.^72^ *Sc. aggregatum* DIA data were acquired in three injections with separate mass ranges (injection 1 = 430–550, 2 = 550–690, 3 = 690–930 m/z) for 120 minutes and *Schizochytrium* sp. 20888 DDA data were acquired in one injection for 60 minutes. Both experiments were performed with HCD at 30% intensity on a Orbitrap Fusion Lumos mass spectrometer (Thermo Scientific).

Gene predictions for nuclear and viral genomes were used as reference databases. DIA raw data were de-multiplexed using msconvert (ProteoWizard, version 3.0, peakPicking = ‘vendor’, demultiplex = overlap only) and analyzed using FragPipe software with DIA-NN (version 1.8.2) for quantification. The three injections intensities were summed for the sample. DDA data were converted with msconvert (ProteoWizard, version 3.0, peakPicking = ‘vendor’) and analyzed in FragPipe using the Label-Free Quantification with Match-Between-Runs (LFQ-MBR) workflow.

## ACKNOWLEDGMENTS

We thank Mary Ann Trevors at the Electron Microscopy Core Facility (Dalhousie University) for preparing samples for transmission electron microscopy, as well as Ping Li at the Scientific Imaging Suite (Dalhousie University) for use of the transmission electron microscope. We also thank Christopher Hughes for extensive support from Dalhousie’s Biological Mass Spectrometry Core Facility. Heroen Verbruggen, Cheong Xin Chan, and Cintia Iha are thanked for support and discussion in the analysis of the *Durusdinium trenchii* genome assemblies. Jackie Collier and Joshua Rest are acknowledged for providing the *Sc. aggregatum* and *T. aureum* cultures used in this study, and Tom Delmont is thanked for feedback on taxonomic assignment of mirusviral sequences. During the preparation of this manuscript, we became aware of a preprint by Yutin et al. (reference 51), who discovered and analyzed a subset of the ‘skiavirus’ sequences analyzed herein. We sincerely thank Eugene Koonin, Nataliya Yutin and their co-authors for helpful discussion of preprinted results.

## DATA AVAILABILITY

Genomic and proteomic data generated in this study are available via figshare at the following link: https://figshare.com/s/abfa66004ba3d7ddfcc4. All other sequence data analyzed herein were retrieved from public data repositories as described in the text.

## EXTENDED DATA LEGENDS

**Extended Data Fig. 1.**
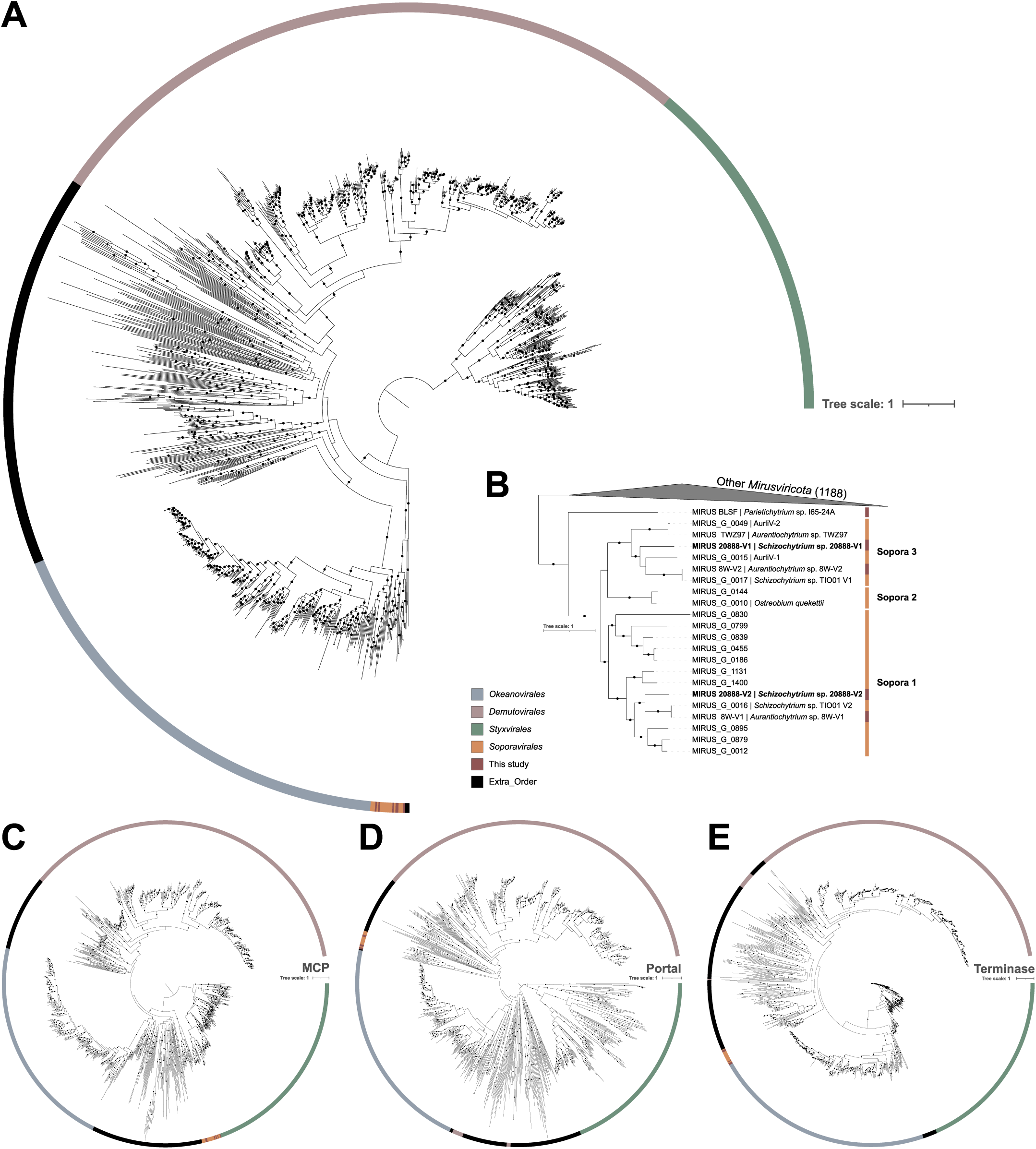
Mirusvirus phylogeny including new mirusviruses from *Schizochytrium* sp. 20888. **a,** IQTree maximum likelihood phylogenetic tree of concatenated MCP-Portal-Terminase proteins (1210 sequences, 2738 amino acid sites, Q.pfam+F+I+R10) and rooted at *Styxvirales*. **b,** Collapsed tree from **a** with *Soporavirales* families annotated. Single gene trees for **c,** MCP (1090 sequences, 1338 amino acid sites, Q.pfam+F+R10) **d,** Portal (994 sequences, 705 amino acid sites, LG+F+R10) and **e,** Terminase (1045 sequences, 695 amino acid sites, LG+F+I+R10). Strong statistical support (aLRT ≥ 80 and UFBoot ≥ 95) is indicated by dots on each branch.

**Extended Data Fig. 2.**
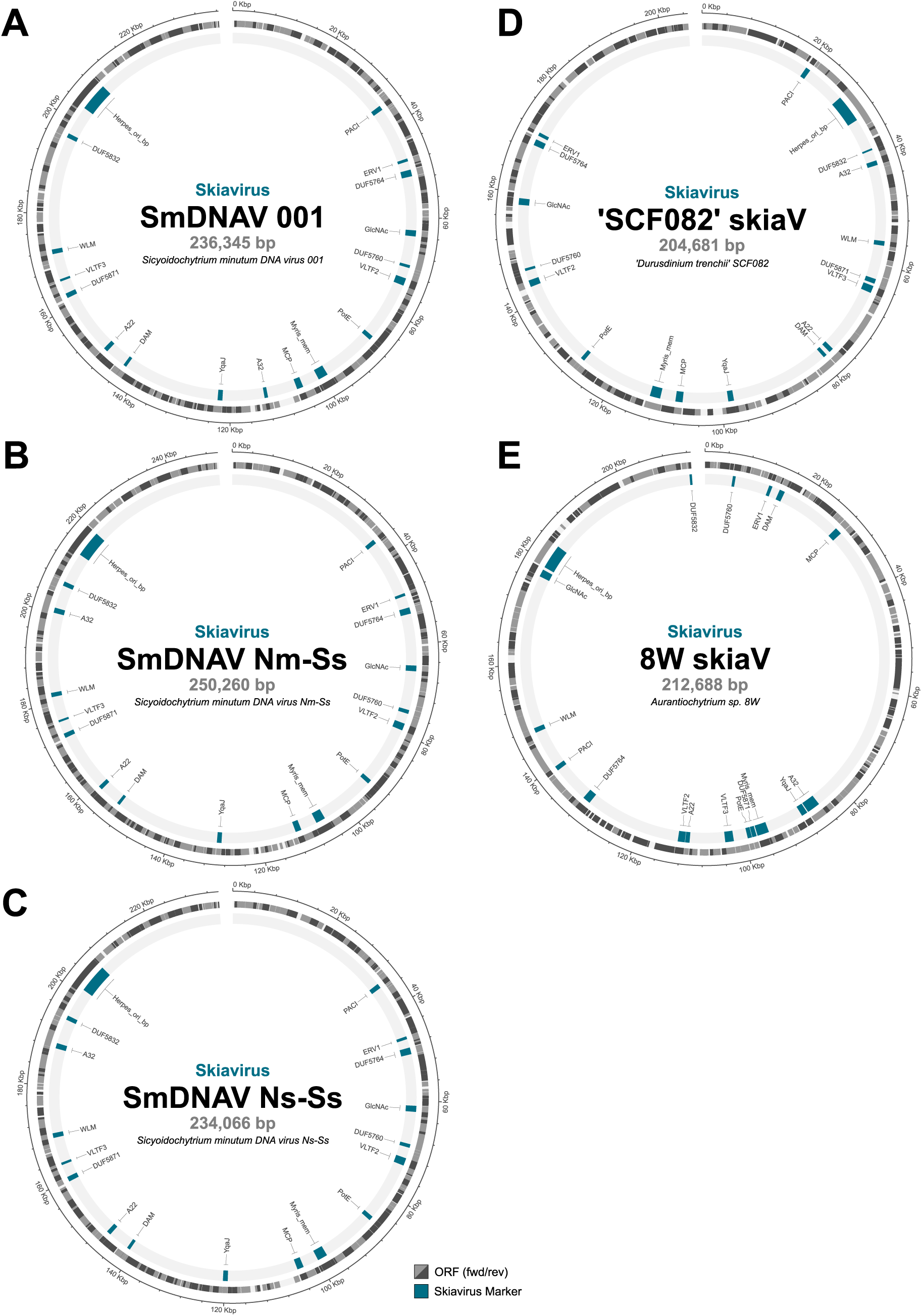
Skiavirus genomes. **a, b,** and **c**, Physical maps of the genomes of *Sicyoidochytrium minutum* DNA viruses 001, Nm-Ss, and Ns-Ss, respectively. Skiaviruses found in additional genome assemblies are also shown as follows: **d,**‘*Durusdinium trenchii’* SCF082 and **e,** *Aurantiochytrium* sp. 8W.

**Extended Data Fig. 3.**
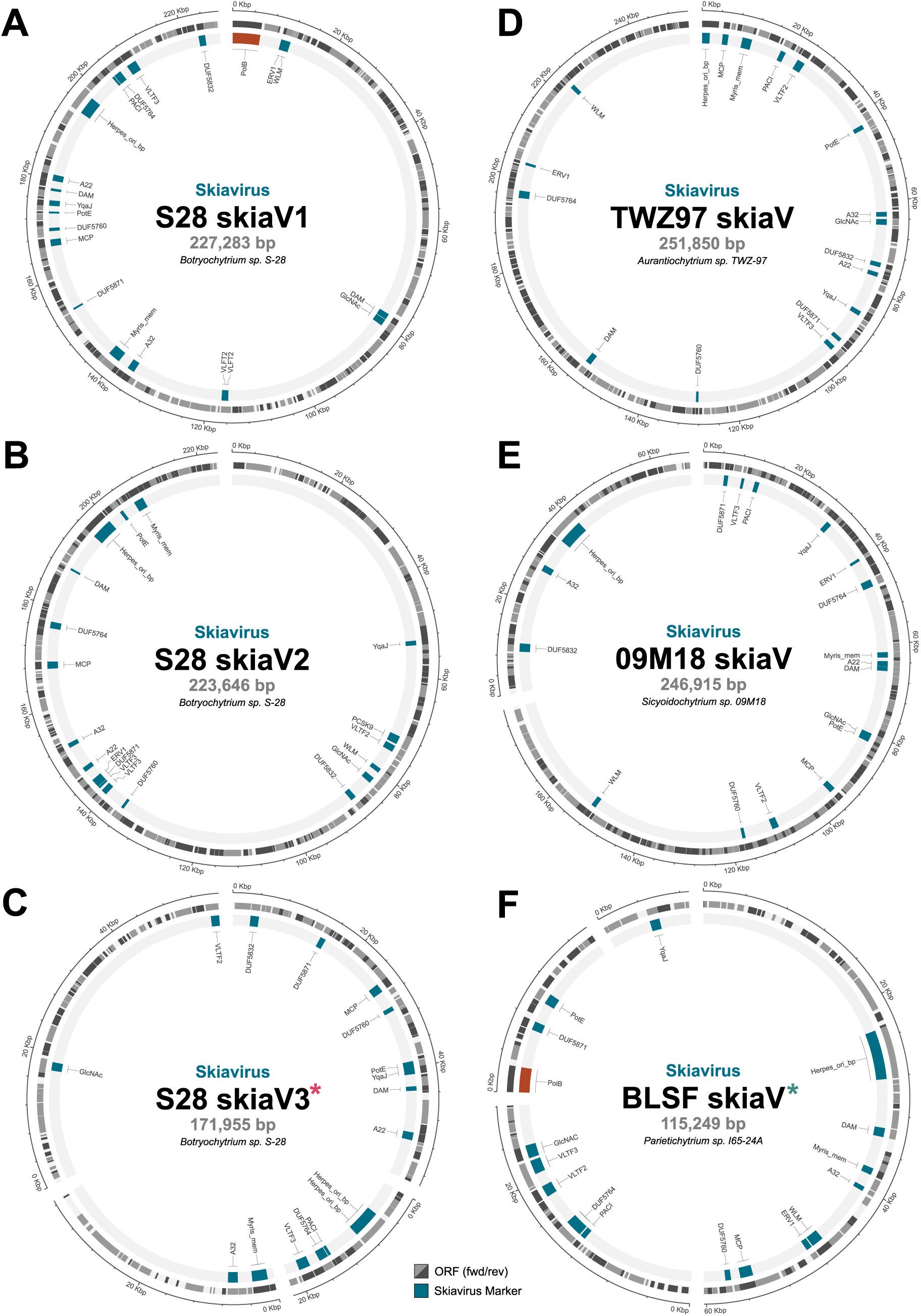
Skiavirus genomes. Physical maps of the genomes of *Botryochytrium* sp. S-28 skiaviruses **a,** S28-V1, **b,** S28-V2 and **c,** S28-V3 respectively. Skiaviruses found in additional genome assemblies are also shown as follows: **d,** *Aurantiochytrium* sp. TWZ-97, **e,** *Sicyoidochytrium* sp. 09M18 and **f,** *Parietichytrium* sp. I65-24A. Asterisks indicate partial skiavirus genomes S28-V3 and BLSF.

**Extended Data Fig. 4.**
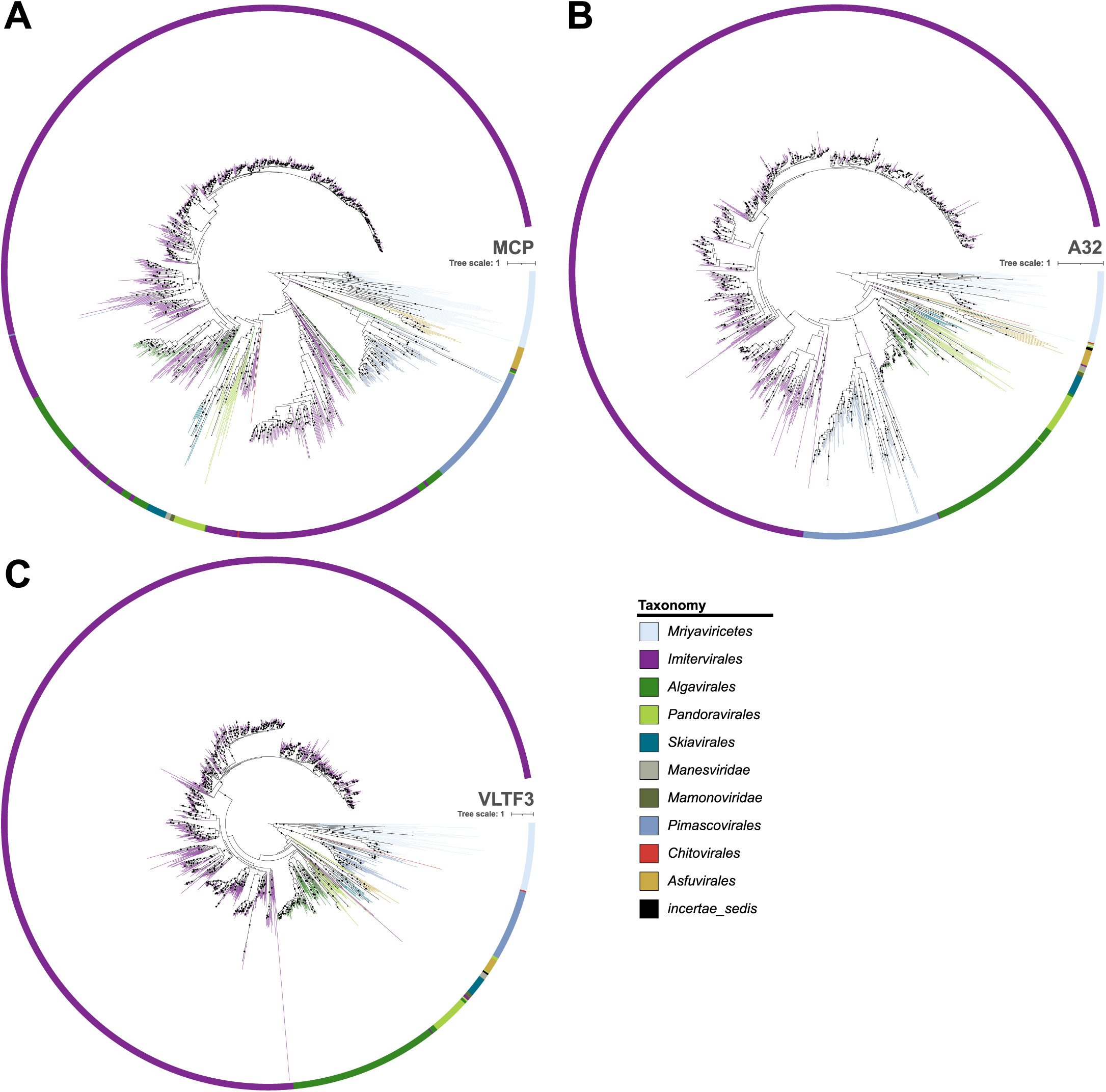
Individual phylogenetic trees for *Nucleocytoviricota* with new proposed order *Skiavirales*. IQTree maximum likelihood phylogenetic trees shown are **a,** MCP (1212 sequences, 640 amino acids sites, LG+F+R10) **b,** A32 (1234 sequences, 297 amino acid sites, model = Q.yeast+R10) and **c,** VLTF3 (1159 sequences, 420 amino acid sites, Q.INSECT+F+I+R10). Strong support (aLRT ≥ 80 and UFBoot ≥ 95) is indicated by dots on each branch. Trees are rooted at *Mriyaviricetes*.

**Extended Data Fig. 5.**
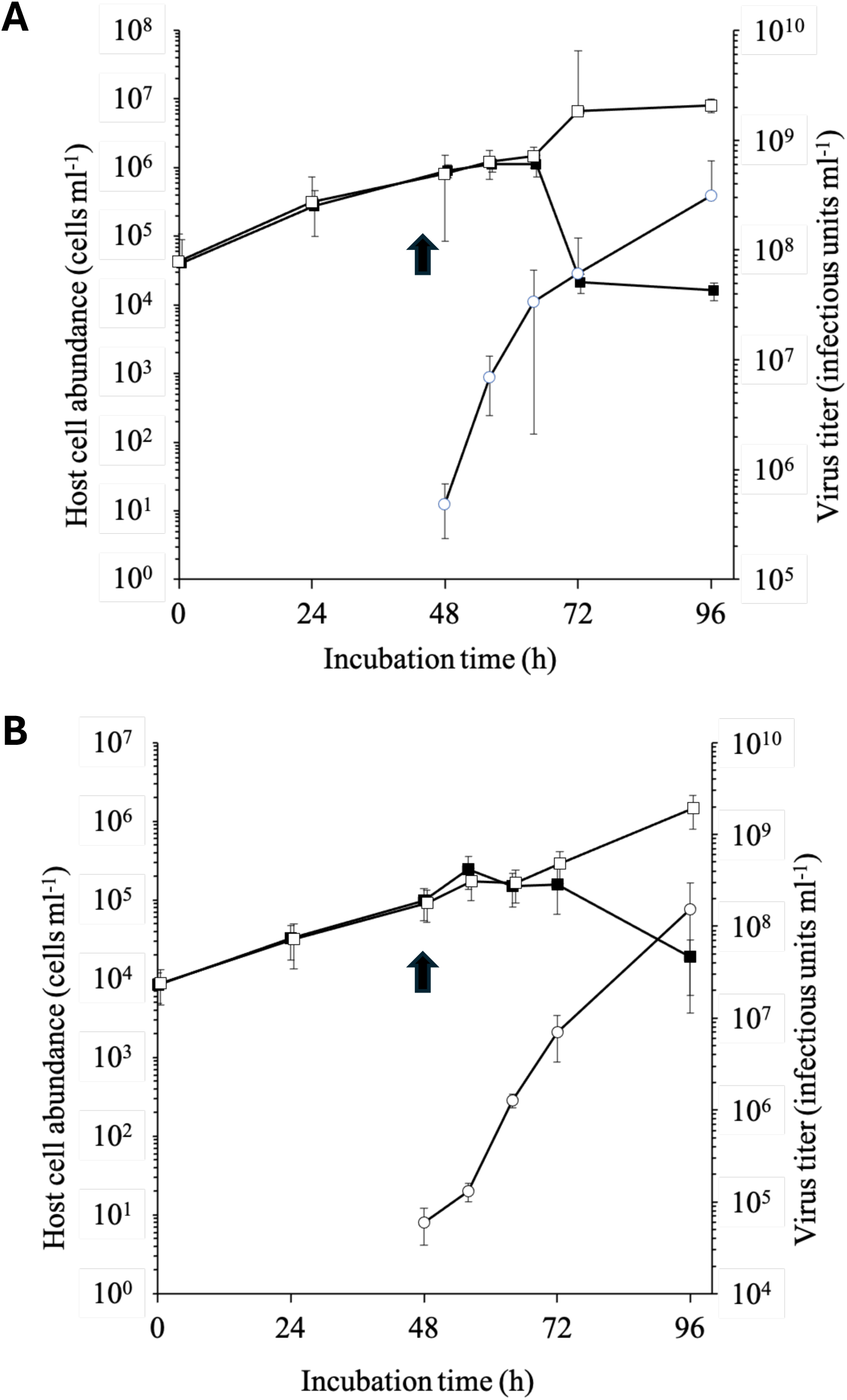
Infectivity of SmDNAV strains on *Sicyoidochytrium* sp. 09M18. Figure shows changes in abundance of 09M18 cells with (closed squares) or without (open squares) viral inoculation, and the viral titer (open circles). Inoculations of SmDNAV strains **a,** Nm-Ss and **b,** Ns-Ss were performed in the exponentially growing phase of host cultures (arrow). Error bars represent standard deviation.

**Extended Data Fig. 6.**
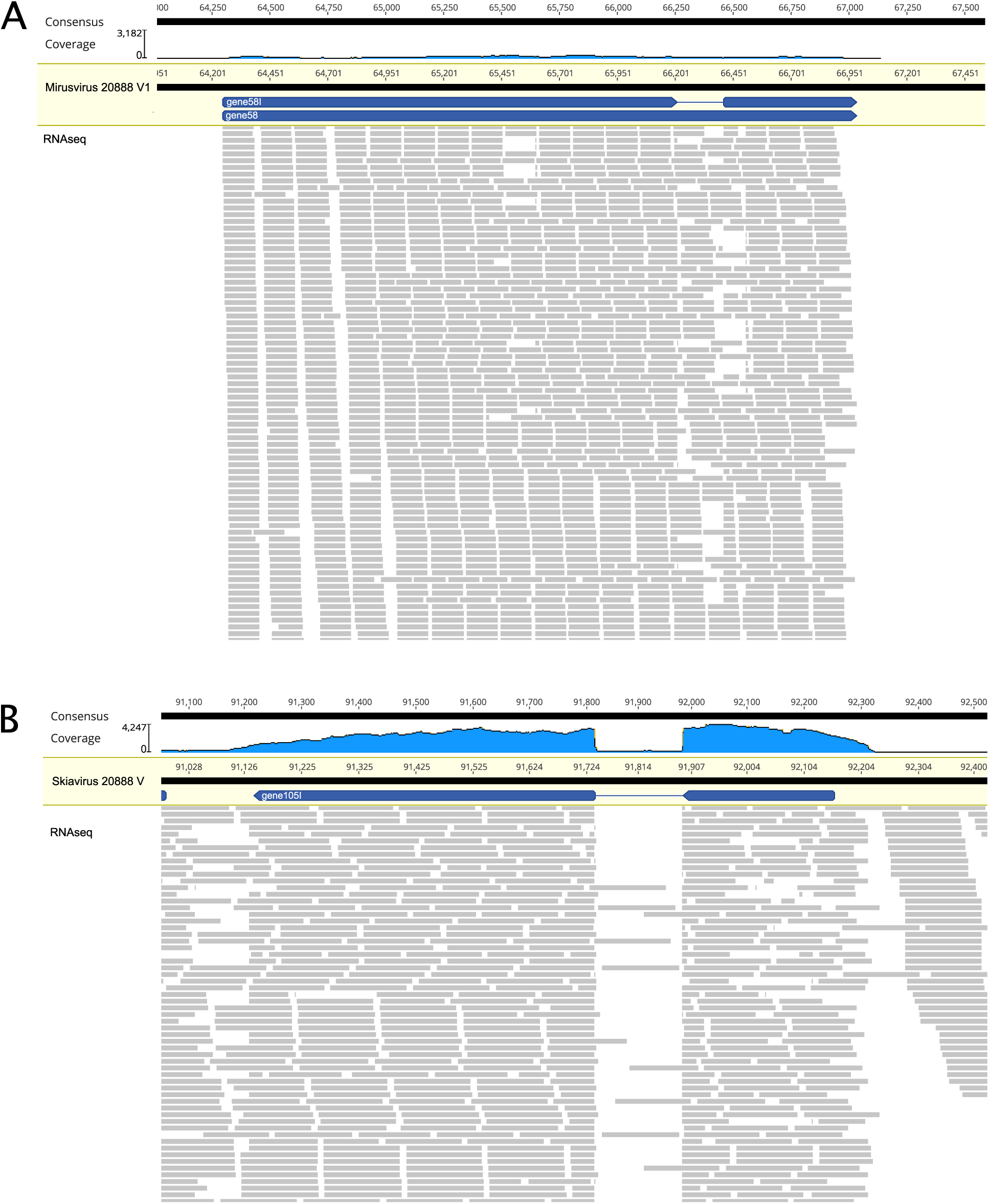
Introns in the viral genomes of *Schizochytrium* sp. 20888. **a,** RNA-seq reads mapped to the *Schizochytrium* sp. 20888-V1 assembly shows low support for a minor alternate isoform of an ankyrin repeat (gene-58 and gene-58I). **b,** a highly expressed 20888-skiaV ORFan (gene-105I) with a strongly supported intron.

**Extended Data Fig. 7.**
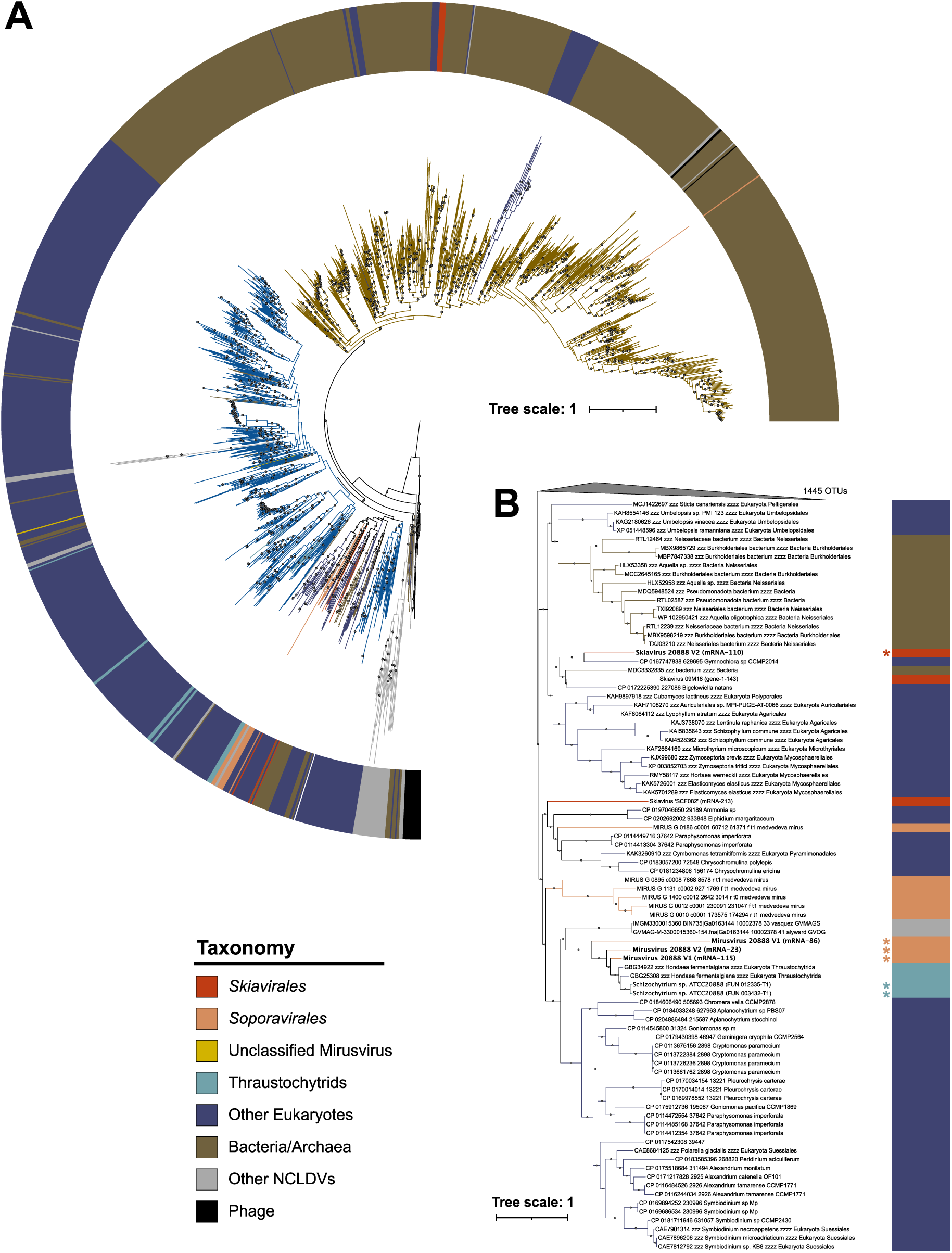
Metallophosphoesterase phylogeny. **a,** Phylogenetic tree of metallophosphoesterase (MPP) with a wide taxonomic distribution. **b,** Collapsed tree from **a** which suggests gene exchange between skiaviruses (red), *Soporavirales* mirusviruses (orange), and thraustochytrid (host) nuclear genomes (teal). Asterisks indicate sequences from *Schizochytrium* sp. 20888.

**Extended Data Fig. 8.**
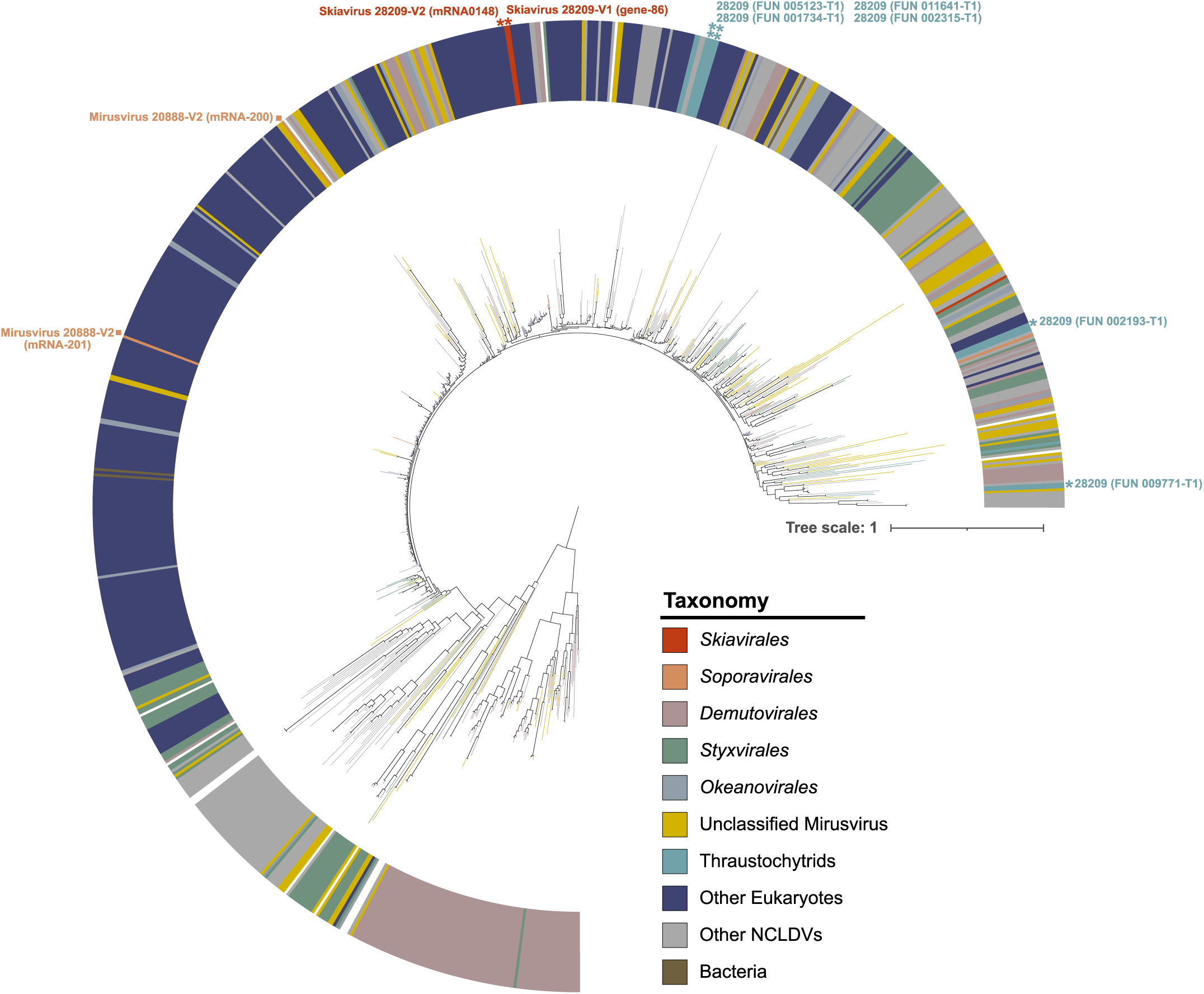
Histone H3 phylogeny. Phylogenetic tree of histone H3 suggests that the histone H3 genes in 28209-skiaV1 and 28209-skiaV2 (red) share ancestry with the nuclear homologs of thraustochytrids (teal) and other eukaryotes (dark blue), with H3 genes in the *Soporavirales* mirusvirus genomes of 20888-V1 and 20888-V2 (orange squares), and with other orders of mirusviruses. Asterisks indicate sequences from *Sc. aggregatum* 28209.

**Extended Data Fig. 9.**
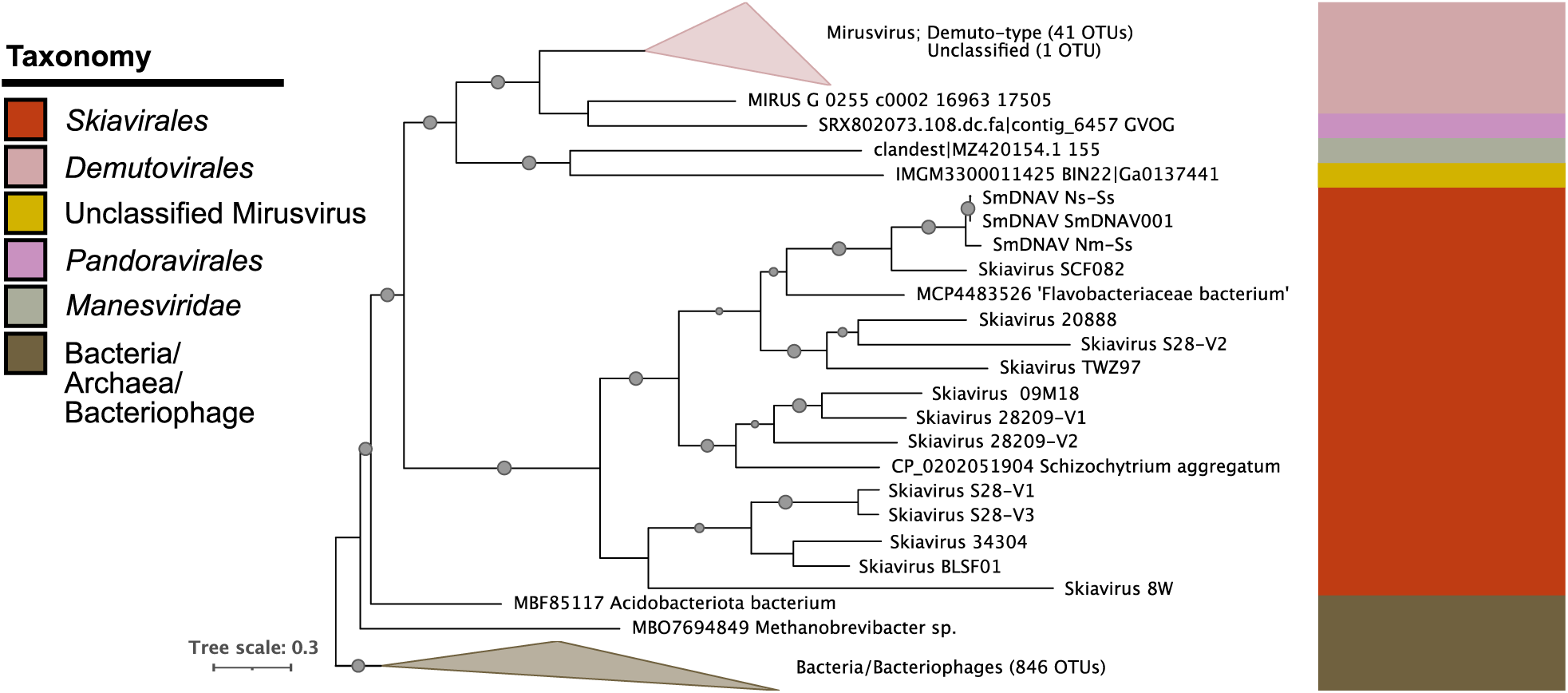
Phylogenetic tree of DNA adenine methyltransferase (DAM), demonstrating gene exchange between skiaviruses (red) and the mirusvirus Order *Demutovirales* (pink).

**Extended Data Table 1.** Nuclear genome summary statistics for long-read sequencing based assemblies of *Schizochytrium* sp. ATCC 20888, *Schizochytrium aggregatum* ATCC 28209 and *Thraustochytrium aureum* ATCC 34304. BUSCO completeness is based on comparison of the genome or predicted protein-coding genes to the stramenopile dataset (v6; stramenopiles_odb12; n=697); C=complete, S=single copy. D=duplicated, F=fragmented, M=missing.

|  | *Schizochytrium* sp.<br>ATCC 20888 | *Schizochytrium*<br>*aggregatum*<br>ATCC 28209 | *Thraustochytrium*<br>*aureum*<br>ATCC 34304 |
| --- | --- | --- | --- |
| Contigs | 30 | 172 | 177 |
| Genome size | 40.09 Mbp | 42.53 Mbp | 39.96 Mbp |
| N50 / L50 | 1.77 Mbp / 10 | 1.33 Mbp / 10 | 0.94 Mbp / 14 |
| Coverage | 53X | 75X | X |
| # N's per 100<br>kbp | 0 | 354 | 0 |
| # of N gaps | 0 | 30 | 0 |
| GC content<br>(%) | 56.65% | 63.4% | 62.9% |
| Repeats | 3.37 Mbp (8.42%) | 6.10 Mbp (14.34%) | 2.72 Mbp (6.81%) |
| Protein-coding genes<br>(CDS) | 13,502 | 13,987 | - |
| BUSCO<br>(genome) | C:96.7%<br>[S:96.1%,D:0.6%],<br>F:1.3%,M:2.0% | C:93.4%<br>[S:91.8%,D:1.6%],<br>F:2.4%,M:4.2% | C:93.0%<br>[S:92.3%,D:0.7%],<br>F:2.7%,M:4.3% |
| BUSCO<br>(CDS) | C:97.4%<br>[S:95.7%,D:1.7%],<br>F:1.1%,M:1.4% | C:92.3%<br>[S:91.1%,D:1.1%],<br>F:4.7%,M:3.0% | - |

**Extended Data Table 2.**
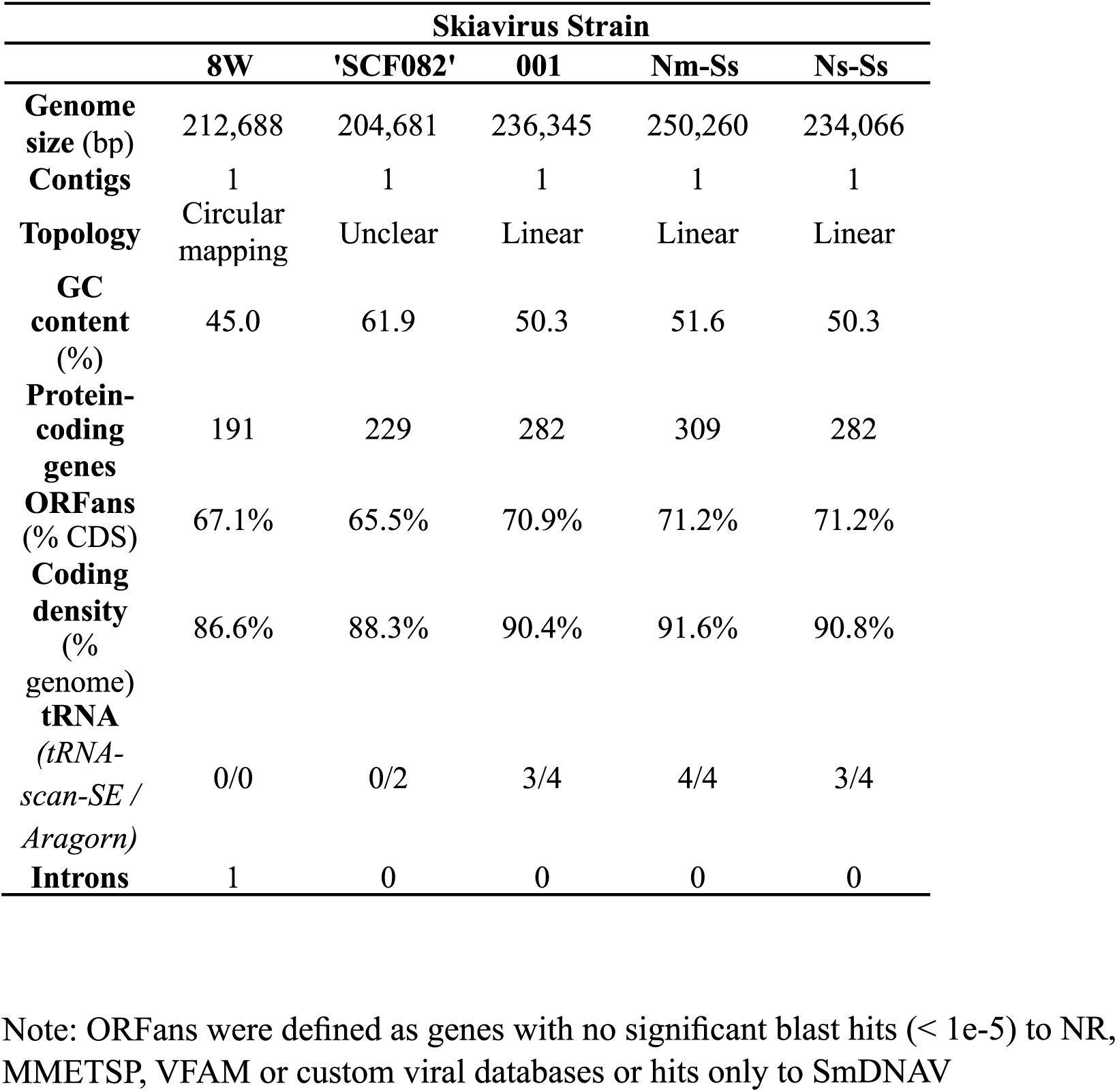
Genome characteristics of five complete skiavirus strains.

**Extended Data Table 3.**
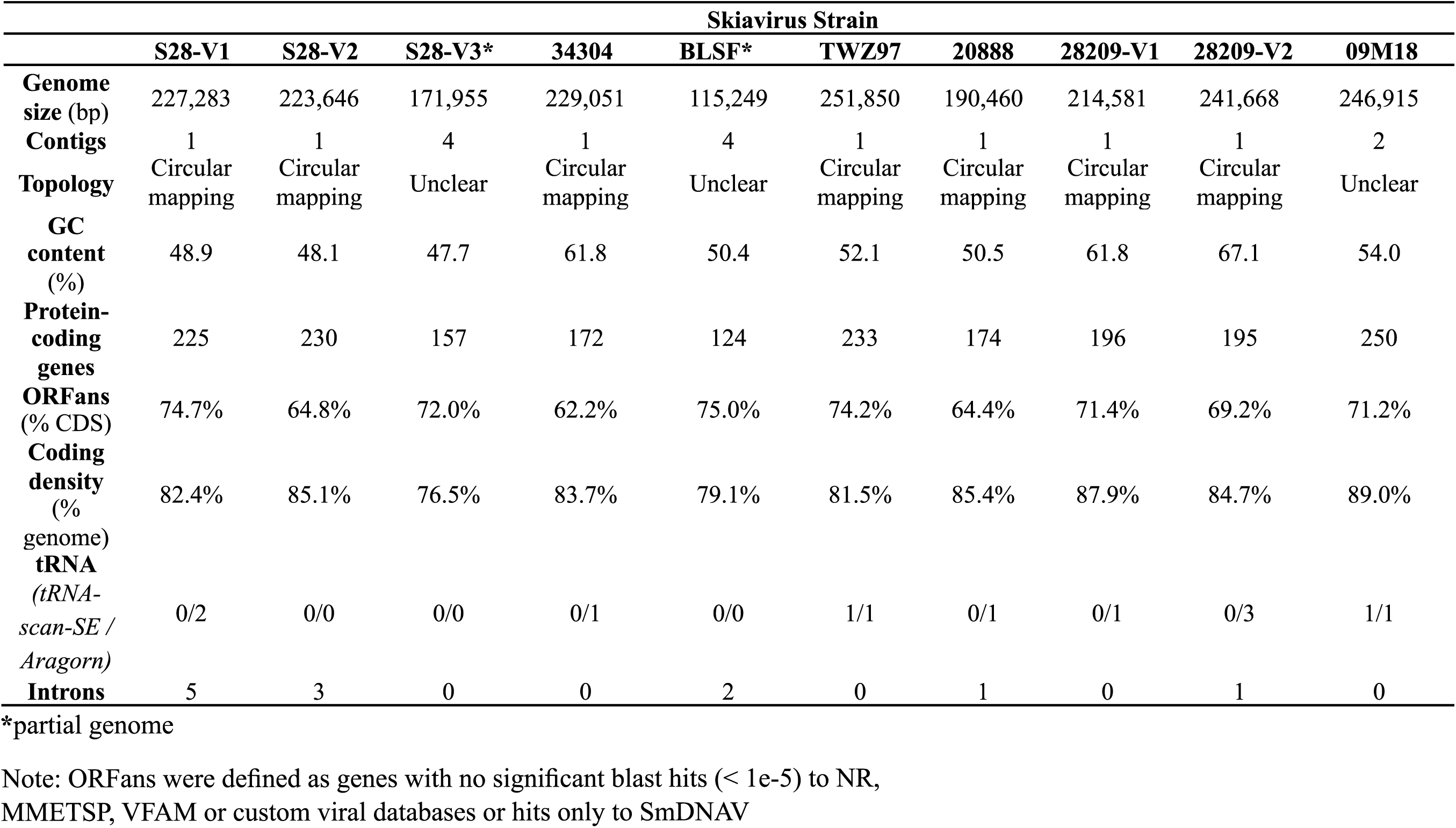
Genome characteristics of 10 complete or near complete skiavirus strains.

**Extended Data Table 4.** Characteristics of two mirusvirus genomes in *Schizochytrium* sp. 20888.

|  | 20888-V1 | 20888-V2 |
| --- | --- | --- |
| Genome size (bp) | 276,422 | 296,200 |
| Contigs | 1 | 1 |
| Topology | Circular mapping | Circular mapping |
| GC content (%) | 64.2 | 52.3 |
| Protein-coding genes | 222 | 212 |
| ORFans<br>(% CDS) | 74.8% | 65.6% |
| Coding density<br>(% genome) | 88.2% | 84.4% |
| Introns | 1 | 0 |

